# Global substrate identification and high throughput *in vitro* dephosphorylation reactions uncover PP1 and PP2A-B55 specificity principles

**DOI:** 10.1101/2023.05.14.540683

**Authors:** Jamin B. Hein, Hieu T. Nguyen, Dimitriya H. Garvanska, Isha Nasa, Yinnian Feng, Blanca Lopez Mendez, Norman Davey, Arminja N Kettenbach, Polly M. Fordyce, Jakob Nilsson

**Affiliations:** Novo Nordisk Foundation Center for Protein Research, Faculty of Health and Medical Sciences, University of Copenhagen, Denmark; Biochemistry and Cell Biology, Geisel School of Medicine at Dartmouth College, Hanover, NH, USA; Department of Bioengineering, Stanford University, Stanford, CA 94305, USA; Division of Cancer Biology, The Institute of Cancer Research, 237 Fulham Road, London SW3 6JB, UK; Department of Genetics, Stanford University, Stanford, CA 94305, USA; Sarafan ChEM-H, Stanford University, Stanford, CA 94305, USA; Chan Zuckerberg Biohub, San Francisco, CA 94110, USA

## Abstract

Phosphoprotein phosphatases (PPPs) dephosphorylate Serine (Ser)/Threonine (Thr) residues to regulate major signaling pathways and cellular transitions. Despite the central role of PPPs the substrates in most cellular processes and the determinants of phosphatase specificity are poorly understood. This is because methods to investigate this at scale are lacking. Here we develop a novel *in vitro* assay, MRBLE:Dephos, that allows multiplexing of dephosphorylation reactions to determine phosphatase preferences. Using MRBLE:Dephos, we establish amino acid preferences of the residues surrounding the dephosphorylation site for PP1 and PP2A- B55, which reveals common and unique preferences for the two phosphatases. To compare the MRBLE:Dephos results to cellular substrates, we focused on mitotic exit that requires extensive dephosphorylation by PP1 and PP2A-B55. We use specific inhibition of PP1 and PP2A-B55 in mitotic exit lysates coupled with quantitative phosphoproteomics to identify more than 2000 regulated phosphorylation sites. Importantly, the sites dephosphorylated during mitotic exit reveal key signatures that are consistent with the MRBLE:Dephos results. We use these insights to specifically alter INCENP dephosphorylation kinetics at mitotic exit, resulting in defective cytokinesis thus underscoring the biological relevance of our determined specificity principles. Finally, we provide a comprehensive characterization of PP1 binding motifs and demostrate how binding of phosphatases to substrates shapes dephosphorylation specificity. Collectively, we develop novel approaches to advance our ability to investigate protein phosphatases and use these to provide a framework for understanding mitotic exit regulation by dephosphorylation.

## Introduction

Cellular signaling by dynamic phosphorylation constitutes a key regulatory mechanism and is vital for cellular function [1]. To understand phosphorylation-dependent signaling we need to understand the principles of kinase and phosphatase specificity. Recent progress has been made in the area of kinase specificity through large scale discovery of active site preferences using peptide arrays [2]. Furthermore, large scale substrate discovery for kinases has been greatly facilitated by the use of specific small molecule inhibitors [3, 4]. In contrast methods that can address protein phosphatase specificity *in vitro* at scale as well as robust substrate identification methods are limited. Establishing such methods is essential for advancing our understanding of dynamic phosphoyraltion and how this is regulated to control cellular signaling.

PP1 and PP2A are members of the phosphoprotein phosphatase (PPP) family and are responsible for the majority of Ser/Thr dephosphorylation in cells [5-7]. PP1 is an evolutionarily conserved phosphatase with three nearly identical isoforms of the catalytic subunit (PP1Cα-γ) [8, 9]. PP1 functionally works as a dimeric holoenzyme consisting of the catalytic subunit bound to a regulatory subunit. Many PP1 regulatory subunits have been identified, and the PP1 catalytic subunit likely does not exist as a monomer. These complexes confer phosphatase activity towards specific substrates by influencing PP1 localization and changing its substrate specificity due to modulation of the PP1 catalytic site [8]. A key mechanism for PP1 holoenzyme formation and substrate interaction is the binding of PP1 to short linear motifs (SLiMs) in proteins. The RVxF motif is the most common SLiM, but several other PP1 binding motifs have been identified (SILK, MyPhone, KiR) [8, 9]. Two recent studies using either a synthetic peptide library approach coupled with a mass spectrometry read-out (PLDMS), or time-resolved mass spectrometry after PP1C depletion started to uncover dephosphorylation site selectivity [10, 11]. These studies revealed a preference for pThr sites and that basic residues surrounding the phosphorylation sites allow for rapid dephosphorylation.

In contrast to PP1, PP2A is an obligatory trimeric holoenzyme with a catalytic subunit (2 isoforms PP2CA/B), a scaffold (2 isoforms PPP2R1A/B), and a regulatory B regulatory subunit (B55, B56, PR72, or Striatin) [6, 7]. Insight into PP2A substrates and specificity has come from different approaches, including the PLDMS approach with PP2A catalytic subunit alone or B55 or B56 depletion/inhibition in cells or mitotic lysates coupled with quantitative mass spectrometry [10, 12, 13]. These studies also highlighted a preference for basic residues N terminal to the phosphorylation site.

Despite the recent insight into PP1 and PP2A specificity, a systematic *in vitro* and *in vivo* comparison of phosphorylation site preference is lacking making it difficult to interpret results obtained from cells or lysates. Here we develop a novel high throughput *in vitro* dephosphorylation assay, MRBLE-Dephos, to provide novel insight into active site preferences for these phosphatases. We establish thousands of PP1 and PP2A-B55 substrates at mitotic exit which confirm several of the key findings from MRBLE-Dephos and provides important insight into mitotic exit regulation by protein phosphatases. Collectively our results provide tools and resource for understanding signal regulation by protein phosphatases and reveals that phosphatases like kinases have active site specificity.

## Results

### A novel high throughput *in vitro* dephosphorylation assay

A systematic analysis of protein phosphatase preference requires the development of a quantitative dephosphorylation assay allowing measurement of multiple dephosphorylation reactions in one experiment. To obtain data at a high throughput scale, we developed a novel multiplexed bead-based dephosphorylation assay (MRBLE:Dephos) (Fig. 1A). MRBLE:Dephos relies on microfluidically-produced hydrogel beads containing ratiometric combinations of lanthanide nanophosphors (MRBLEs), each of which forms a unique spectral code that can be identified via imaging alone [14-16]. In prior work, we demonstrated that peptides can be chemically synthesized directly on MRBLEs, making it possible to quantify binding of a fluorescently-labeled protein to 96 bead-bound peptides in parallel, using only small amounts of reagents [17, 18]. We have now extended this assay to quantify protein-dependent dephosphorylation by synthesizing 94 phosphorylated Ser or Thr containing peptides with basic, acidic, or hydrophobic amino acids surrounding the phosphorylation site on MRBLEs with a 1:1 linkage between the identity of the synthesized peptide and the embedded spectral code (Fig. 1A). In this set of 94 peptides, we varied key positions (X) in a core sequence, LGAXXXSp/TpXLXXVSA, to test the effect of acidic, basic or hydrophobic residues on PP1γ and PP2A-B55α dephosphorylation kinetics (for simplicity referred to as PP1 and PP2A-B55). The peptide set covered the same sequences with either a phosphorylated serine or threonine and half of the peptides contained a proline residue right after the phosphorylation site. After incubation with phosphatase for different amounts of time (15, 30, 60, 120, 240 minutes), we incubated MRBLE-bound peptides with the pIMAGO reagent [19, 20] (which binds specifically to phosphorylated proteins and has a biotin moiety) and DyLight 650-labeled streptavidin (which binds the pIMAGO biotin moiety). After washing, imaging of MRBLEs in both the lanthanide and fluorescence channels identifies embedded codes (and thus the peptide associated with each bead) and quantifies DyLight 650 fluorescence (and thus the amount of phosphate remaining). Comparing observed fluorescence with negative control samples not exposed to phosphatases then makes it possible to follow phosphatase-dependent dephosphorylation of 94 peptides over time within a single small volume. To test the performance of the assay, we used Lambda phosphatase to dephosphorylate all 94 peptides on the mixed beads (Fig. 1B). Fluorescence images of MRBLEs post-phosphatase treatment showed a loss of DyLight 650 fluorescence, confirming the ability to use this assay to detect dephosphorylation (Fig. 1B).

**Figure 1.**
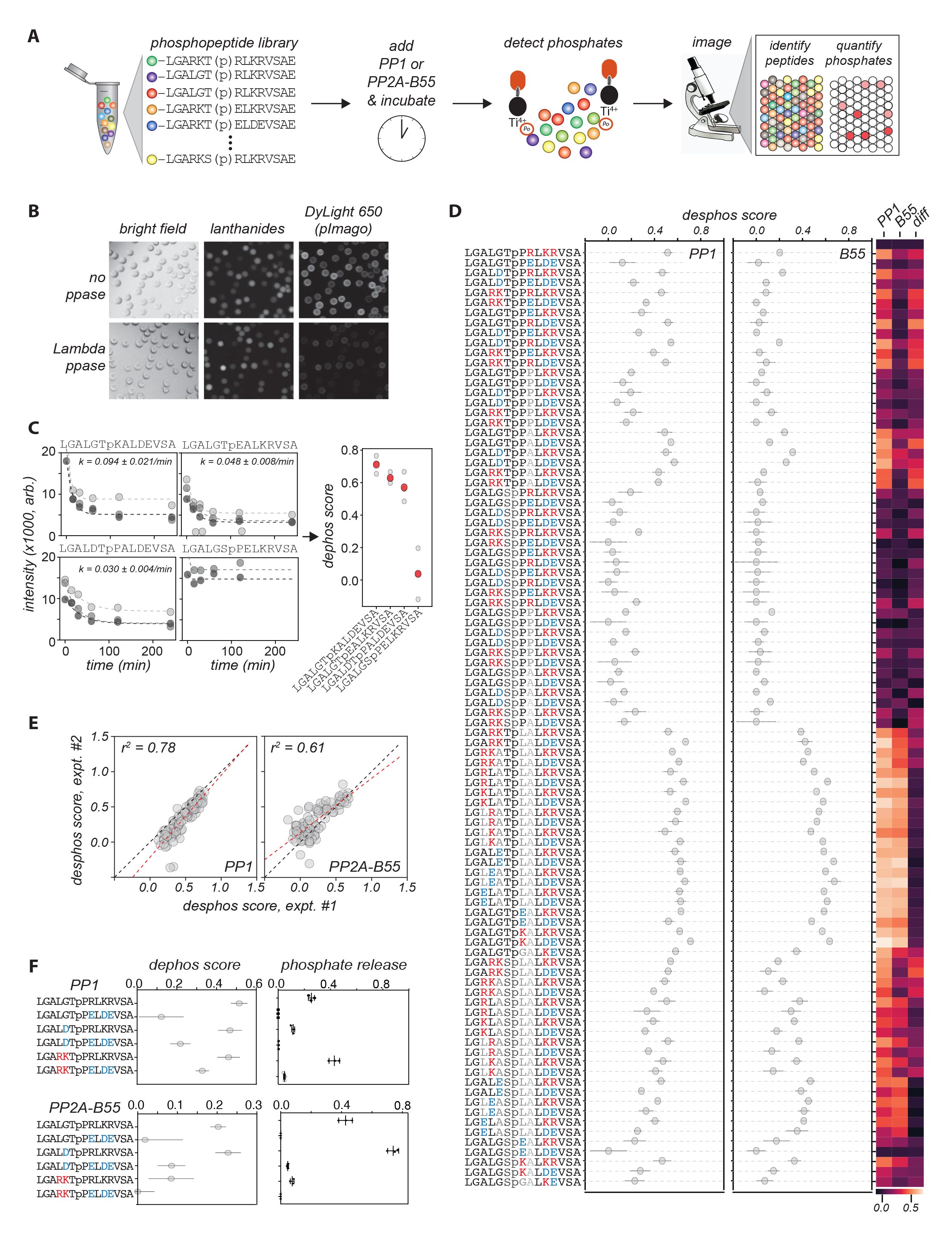
MRBLE:Dephos - a novel multiplex *in vitro* dephosphorylation assay. **A**) Schematic of the MRBLE:Dephos assay workflow. Phosphorylated peptides are chemically synthesized on MRBLEs spectrally encoded beads with a 1:1 linkage between peptide sequence and embedded code. After treatment with protein phosphatases, MRBLE-peptide libraries are incubated with biotinylated pImago and DyLight 650 streptavidin, washed, and imaged to identify embedded spectral codes and quantify bead-bound fluorescence. **B**) Example images of beads either untreated or treated with Lambda phosphatase via bright field imaging, imaging in a lanthanide channel, and imaging in the DyLight 650 channels; DyLight 650 reports on the phosphorylation status of the peptides. **C**) Measured DyLight 650 fluorescence intensities after various incubation times for PP1 interacting with 4 example peptides. Marker values represent the median intensity across all beads measured at each time point. Each marker shade indicates a different experimental replicate, dashed lines indicate single exponential fits, and the annotation denotes the mean fitted rate across experiments (error is given as the standard deviation across the 3 fits). **D**) Dephosphorylation scores across 94 peptides for PP1 (left) and B55 (right); markers denote median values across 3 experimental replicates and error bars denote standard deviations. Heat maps at right show mean dephosphorylation scores for PP1 and B55 and computed differences in scores between proteins. **E)** Scatter plots comparing per-peptide measured dephosphorylation scores across 2 replicate experiments for PP1 (left) and B55 (right); black dashed line indicates 1:1 line and red dashed line indicates linear regression with annotated Pearson correlation coefficient. **F)** In vitro phosphatase assays with indicated peptides comparing results of MRBLE-dephos to dephosphorylation assays using malachite green.

We next monitored dephosphorylation of the 94 peptides after incubation with purified recombinant PP1 and PP2A-B55 affinity purified from HeLa cells (Fig. 1C-E). Consistent with sequence-dependent differences in dephosphorylation, DyLight 650 fluorescence signals exhibited a variety of behaviors over time, from decreasing rapidly to remaining relatively constant; for some peptides, observed time-dependent changes were well-fit by a single exponential (Figs. 1C, S1-2). For each peptide sequence, we calculated a ‘dephosphorylation score’ by: (1) dividing the median ending fluorescence intensity for all beads (averaged across the last 3 timepoints) by the median starting fluorescence intensity for all beads and then (2) subtracting this normalized quantity from 1 such that higher values correspond to more dephosphorylation (Fig. 1C). Although the PP1 preparation had higher activity than the PP2A-B55 preparation, resulting in much larger differences between initial and final peptide phosphorylation levels for PP1 (Fig. S1-2), ‘dephosphorylation scores’ calculated in this manner were highly reproducible across experimental replicates for both proteins (Figs. 1E, S3-4).

To test if sequence-specific dephosphorylation preferences detected by MRBLE:Dephos are consistent with established phosphatase assays, we compared MRBLE:Dephos results to a standard phosphate release assay with a panel of 6 phosphorylated proline-directed Thr sites and found that MRBLE:Dephos results gave comparable patterns of specificity (Fig. 1F). More broadly, MRBLE:Dephos scores for peptides containing identical sequences outside of the phosphosite for both PP1 and PP2A-B55 revealed a strong preference for phosphorylated threonines (Figs. 1D, 2A). Further classifying peptides by whether they contained a proline at the +1 position shows that proline-directed sites are generally less efficiently dephosphorylated than non-proline directed sites, consistent with previous reports (Figs. 1C, 1D, 2B) [21].

**Figure 2.**
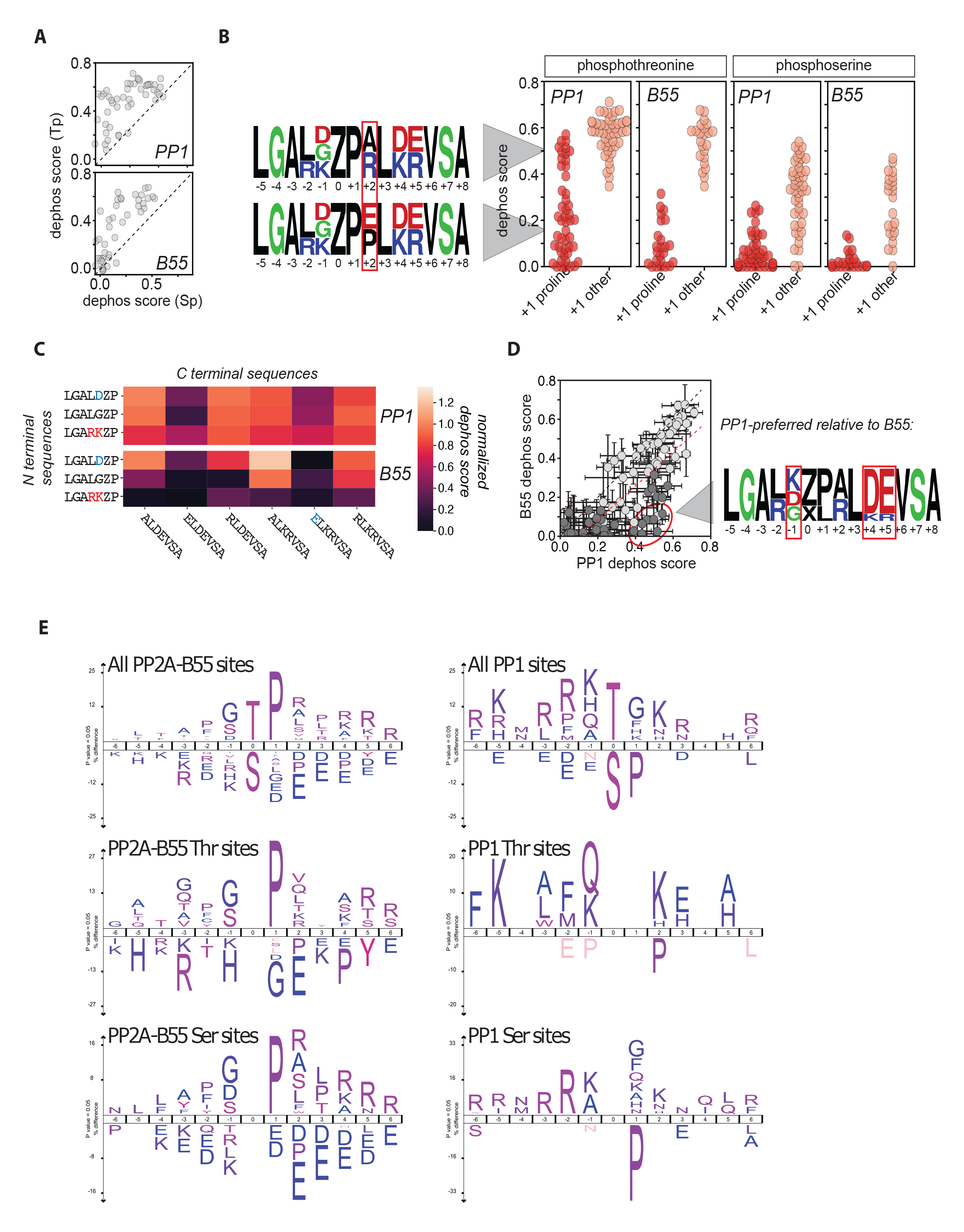
Mitotic exit dephosphorylomes for PP1 and PP2A-B55. **A**) Comparison of dephosphorylation scores for peptides containing identical sequences around phosphothreonine vs phosphoserine sites for PP1 (top) and B55 (bottom); dashed black line indicates 1:1 line. **B**) Dephosphorylation scores for PP1- and B55-mediated dephosphorylation of peptides containing either phosphothreonine (left) or phosphoserine (right) with either a proline (dark red) or other residue (light red) in the +1 position. Weblogos for peptides with higher (top) or lower (bottom) dephosphorylation scores show differential effects of A, R, E, and P in the +2 position; Z and X denote phosphothreonine and phosphoserine, respectively. **C**) Heat maps showing dephosphorylation scores for PP1 (top) and B55 (bottom) for phosphothreonine peptides with a +1 proline and different combinations of N-terminal (rows) and C-terminal (columns) sequences. **D**) Scatter plot comparing B55 *vs.* PP1 dephosphorylation for peptides with the same sequence. Weblogo for sequences that are preferentially dephosphorylated by PP1 show a strong enrichment for D and E at the +4 and +5 positions and a K at the −1 position. **E**) Icelogos for PP1 and PP2A-B55 based on single fully localized phosphorylation sites detected in the screen.

Focusing on proline-directed sites, a common trait for both PP1 and PP2A-B55 is the negative impact of acidic residues C-terminal to the phosphorylation site, in particular position +2 (Fig 2B-D). For residues N-terminal to the phosphorylation site, PP1 overall preferred basic residues in pos −1 and −2 while an acidic or glycine residue at −1 was not well tolerated (Fig. 2C). For PP1 N-terminal basic residues could compensate for the negative impact of an acidic or glycine residue in pos +2 (Fig. 2C). By contrast, PP2A-B55 prefers aspartic acid and glycine residues in −1 while basic residues at −1 and −2 are disliked. However, the presence of an aspartic acid or glycine residue could not compensate efficiently for an acidic residue in pos +2 (Fig. 2C). Relative to PP2A-B55, PP1 displays higher active towards peptides with positive residues before and acidic residues after the phosphorylation sites (Fig. 2D).

In conclusion, we develop a generally applicable high throughput dephosphorylation assay that allows quantitative mapping of protein phosphatase dephosphorylation motifs using only very small amounts of material. We use this to show the existence of optimal dephosphorylation site signatures for PP1 and PP2A-B55, providing a set of rules for accessing whether a phosphorylation site is optimal for one of these phosphatases.

### Dephosphorylomes for PP1 and PP2A-B55 during mitotic exit

To benchmark our *in vitro* data with *in vivo* substrates, we set out to establish “dephosphorylomes” for PP1 and PP2A-B55. We focused on mitotic exit as this cellular transition requires massive dephosphorylation of mitotic kinase sites by PP1 and PP2A-B55. We recently developed an assay for unbiased identification of phosphatase substrates in mitotic cell lysates (Fig. S5) [12, 22]. Nocodazole-arrested cells were lysed in the presence of specific PP1 or PP2A-B55 phosphatase inhibitors. As ATP is quickly depleted in the extract this effectively stops all kinase activity and thus resembles a state of mitotic exit where phosphatases are reversing the action of mitotic kinases. We used the central domain of Nipp1 and thiophosphorylated Arpp19 (thio-Arpp19) as specific natural inhibitors of PP1 and PP2A-B55, respectively [23-25]. Nipp1 prevents binding of RVxF-containing proteins to PP1, in this way preventing holoenzyme formation and thus PP1 dephosphorylation. In contrast, thio-Arpp19 blocks the active site of PP2A-B55, effectively blocking activity. As controls, we used Nipp1 with a mutated RVxF motif preventing PP1 binding and Arpp19 S62A that cannot inhibit PP2A-B55. An antibody detecting all proline-directed threonine phosphorylation (TpP) as well as H3S10 confirmed the specific inhibition of phosphatases upon addition of inhibitors (Fig. S5). We used this setup for unbiased identification of phosphorylation sites that are stabilized after treatment of inhibitors using TMT labelling and phospho-peptide enrichment followed by mass spectrometry. This resulted in the identification of 38,173 phosphorylation sites of which 2116 and 751 were significantly increased in phosphorylation by 1.5-fold or more after PP2A-B55 or PP1 inhibition, respectively (Supp. Table 1). Only 152 sites (5,6%) showed an increase in both conditions attesting to high specificity of the phosphatases. We identified known B55-regulated sites on the B55 inhibitor ENSA S67 and Arpp19 S62 [26], confirming that PP2A-B55 is active in our assay conditions and inhibited when adding thio-Arpp19. Furthermore, the reported PP1-regulated site H3S29 was stabilized specifically by adding Nipp1 [27]. Collectively, these data validated our approach.

Comparison to previous screens of PP2A-B55 substrates revealed an overlap of 124 of 203 phosphorylation sites on 95 proteins identified by Cundell et al [13] as high confidence B55 substrates using siRNA depletion of B55 or MASTL. This 61% overlap between two independently performed phosphoproteomic studies using different approaches is very high, strongly supporting the notion that these are high confidence PP2A-B55 substrates. Surprisingly, for our PP1-dependent phosphorylation sites, we only identified a few sites that were previously shown to be PP1-dependent in MS analysis of mitotic cells depleted of PP1 [11] (13 out of 302) or in phosphatases inhibited in cell lysates after the addition of PP1 catalytic subunit [10] (10 out of 777). The difference in experimental approach and cell cycle stage of cell lysates likely explain this limited overlap.

Closer examination of the increased phosphorylation sites revealed that 1185 of the 2116 sites significantly increased upon Arpp19 addition were detected in the Nipp1 treated sample, and conversely, 548 of the 751 phosphorylation sites significantly increased upon Nipp1 addition were detected in the Arpp19 treated samples. It is likely that not all sites are detected to the same extent because upon inhibition of PP2A-B55 or PP1, phosphorylation site occupancy dramatically increases, thereby enabling the detection of low abundant sites that are otherwise undetectable via LC-MS analysis. Interestingly, proteins like PRC1 were dephosphorylated by both PP1 and PP2A-B55 but at distinct sites, showing that the two phosphatases can act on the same protein potentially to regulate different functional aspects.

For further analyses of phosphorylation preferences of PP2A-B55 and PP1 in mitotic cell lysates, we focused on single, fully-localized phosphorylation sites that were identified in both datasets and uniquely matched one protein. Sites were clustered based on their changes in abundance upon PP2A-B55 and PP1 inhibition, and clusters were assigned as PP2A-B55- or PP1- specific if the sites were significantly regulated by only one of the phosphatases or ambiguous if they were significantly changed in abundance in both datasets. This yielded a phosphoacceptor distribution of Ser:Thr of 65%:35% and 58%:42% for PP2A-B55 and PP1, respectively. Amino acid sequences corresponding to regulated phosphorylation sites and their surrounding residues were entered into Ice Logo using non-regulated phosphorylation sites as background (Fig. 2E). This revealed a strong preference for PP2A-B55 for proline-directed Ser and Thr phosphorylation sites. For PP1, we identified a preference for basic amino acids in the −2 and −3 positions and a deselection of prolines in the +1 position of Ser phosphorylation site. Interestingly, PP1 dephosphorylated Thr phosphorylation sites with any amino acids in the +1 position. Importantly, our cell lysate data are in overall good agreement with our MRBLE:Dephos data on PP1 in terms of a preference for upstream basic residues as well as in the +2 position and deselection of Pro residues in +1. In addition, PP2A-B55 dephosphorylation motifs with glycine or aspartic acid residues in −1 were enriched in our lysate data while we detected deselection of acidic amino acids in +2 for PP2A-B55 in line with the MRBLE:Dephos data. In contrast, the lysate data revealed a strong enrichment of a Pro residue in +1 for PP2A-B55 which our MRBLE:Dephos data showed had a negative impact on dephosphorylation. The exact reason for this is unclear but could suggest that additional factors such as the cis/trans configuration of the +1 Pro residue, which we do not control in MRBLE:Dephos assay, might facilitate efficient PP2A-B55 dephosphorylation *in vivo*. Furthermore, binding of PP2A-B55 to specific interactors resulting in local high concentrations could facilitate dephosphorylation of TP and SP sites.

Collectively our two distinct experimental approaches establish overall consistent results regarding PP1 and PP2A-B55 preferences for dephosphorylation sites and our dephosphorylomes provides novel insights into mitotic exit coordination by PP1 and PP2A-B55.

### Phosphorylation site preference influence dephosphorylation and cell division

To investigate if the uncovered preferences for phosphatases are biologically relevant for mitotic exit, we used INCENP T59 dephosphorylation by PP2A-B55 as a model [28]. During anaphase, the chromosomal passenger complex (CPC) translocates from chromatin to the central spindle. INCENP T59, a Cdk1 phosphorylation site, must be dephosphorylated to ensure timely translocation and accurate cytokinesis (Fig. 3A) [29]. Based on the rules we had established for PP2A-B55, we predicted that changing the residues surrounding T59 would affect dephosphorylation in cells. Our results suggested that substituting the +2 position to glutamic acid (S61E) should have a profound effect on PP2A-B55 dephosphorylation. Entirely consistent with this, PP2A-B55’s ability to dephosphorylate an INCENP^53-67^ S61E peptide was severely impaired, while this mutation only had a small effect on PP1 dephosphorylation likely because of the compensation from the K in −1 (Fig 3B). Importantly, the ability of Cdk1 to phosphorylate T59 was not affected by the S61E mutation (Fig 3C). Using live-cell microscopy, we followed the translocation of Venus-tagged INCENP WT or S61E. In contrast to INCENP WT, S61E did not translocate to the central spindle during anaphase in agreement with a lack of T59 dephosphorylation (Fig 3D). We next depleted endogenous INCENP by RNAi and complemented cells with RNAi resistant Venus-INCENP WT or S61E and followed mitotic progression. Consistent with the inability of INCENP S61E to translocate at mitotic exit, we observed around 60% of binucleated cells in INCENP S61E complemented cells (Fig 3E).

**Figure 3.**
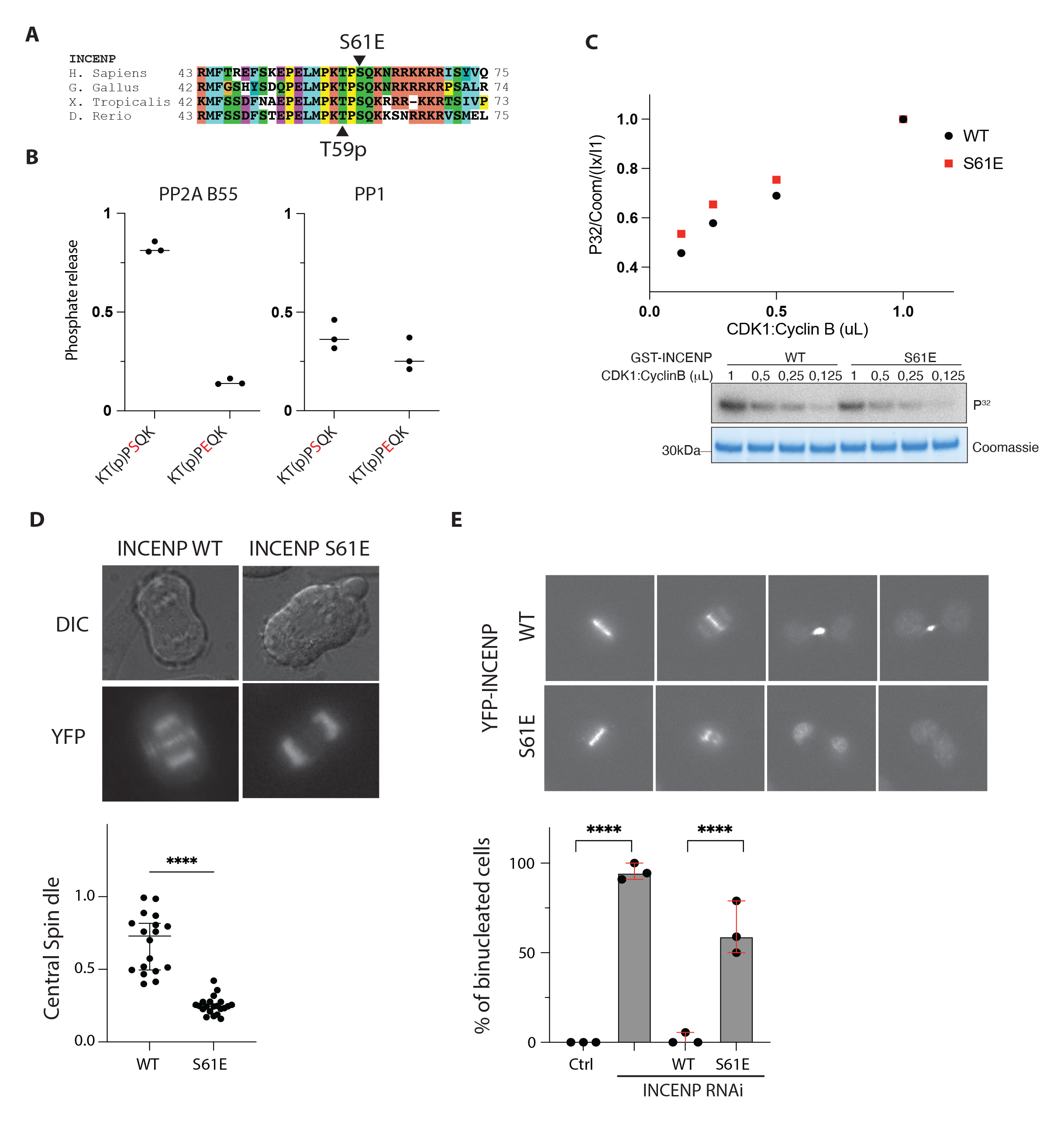
Engineering PP2A-B55 dephosphorylation of INCENP T59. **A)** Sequence conservation around INCENP T59. **B**) In vitro dephosphorylation assays of the indicated peptides with PP2A-B55 and PP1. **C**) In vitro kinase assays with Cyclin B1-Cdk1 and GST-INCENP 45-75 proteins. Representative of 3 independent experiments. **D**) Representative images of YFP-INCENP proteins at anaphase and quantification of central spindle. **E**) Cells depleted of INCENP and expressing the indicated YFP-INCENP constructs where recorded by time-lapse and mitotic progression followed. The percentage of cells becoming binucleated is indicated.

Collectively, we show that the specificity rules we have established are important for regulating biological processes.

### Mitotic phosphatase holoenzymes and orchestration of dephosphorylation

In addition to active site preferences, binding of protein phosphatases to specific regulatory proteins contribute to site-specific dephosphorylation. In the case of PP1, this phosphatase is always in complex with additional proteins and never acts as a free catalytic subunit. To explore this concept in more detail in the context of our mitotic dephosphorylomes, we focused first on defining mitotic interactors for PP1 and PP2A-B55.

We defined interactomes of PP1γ and B55α during mitosis using affinity purification of YFP-tagged subunits and also proximity-dependent ligation using miniTurbo to capture interactions at mitotic exit (Fig. S6) [30, 31]. This resulted in the identification of 923 PP1 interactors and 141 B55 interactors (Supplemental Table 2). Comparing this to the phosphoproteomic data revealed that there were 319 PP1 regulated sites in the PP1 interactome and 100 PP2A-B55 regulated sites in the B55 interactome (Fig. 4A-B). We then integrated these interactomes and information from the STRING database [32] with our phosphorylation site to generate a network of complexes and dephosphorylation sites (Figs. 4C and S7A). For PP1, we also integrated information on known and predicted RVxF motifs [33]. Our networks revealed that both PP1 and PP2A-B55 dephosphorylate a diverse set of proteins regulating core biological processes involved in mitotic exit as well as interphase functions. Furthermore, we observed for PP1 numerous substrates linked to nucleolar function consistent with the localization of PP1 to this structure.

**Figure 4.**
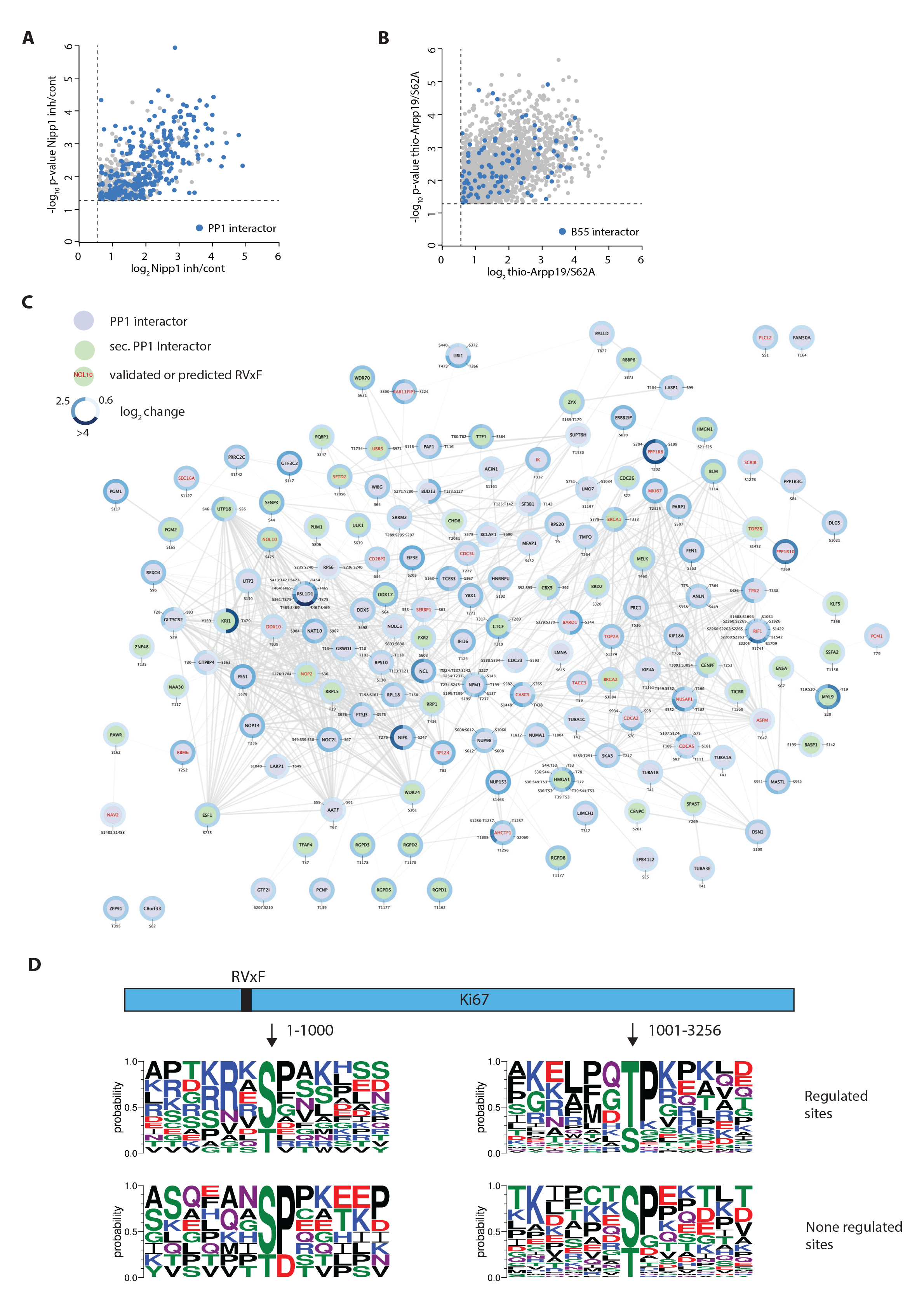
A mitotic exit PP1 dephosphorylation network. **A**) Sites being dephosphorylated by PP1 with sites in a protein identified in interactome approaches indicated by color. **B**) Network of PP1 dephosphorylations indicating proteins containing RVxF motifs and sites regulated in our lysate screen. **C**) Analysis of PP1 regulated sites in Ki67 and the role of the RVxF motif.

Our integration of interactors and regulated sites then allowed us to explore how the binding of phosphatases to proteins might affect dephosphorylation preferences. We first focused on PP1, as RVxF motifs were present in many of our interactors and this allowed us to position PP1 binding precisely in relation to regulated sites in the primary structure of substrates. We used the RVxF-containing protein KI67 as a model since we identified 195 phosphorylation sites on KI67, more than 116 of which exhibited a fold-change >1.5 after PP1 inhibition. KI67 is largely unstructured and spans 3256 amino acids with one RVxF motif positioned at 504-508, and we mapped regulated sites throughout the protein. We generated dephosphorylation motifs for all localized, single phosphorylation sites around the RVxF motif (aa 1-1000) and further away (aa >1000). Sites were clustered into PP1-regulated sites (fold-change of 2) or sites that were not stabilized after PP1 inhibition (Fig 4D). PP1-regulated phosphorylation sites close to the RVxF motif were dominated by phosphorylated Serine (pSer) residues surrounded by basic residues, while more distant sites were enriched for phosphorylated threonine. A higher local concentration of PP1 around the RVxF motif might allow for efficient dephosphorylation of pSer, while pThr can be efficiently dephosphorylated by lower concentrations of PP1. Analyzing the phosphorylation sites on proteins with or without predicted RVxF motif showed a similar pattern as in KI67: dephosphorylation sites on proteins without RVxF were more basophilic and pSer sites were less frequent (Fig. S8). A similar tendency was observed when we compared regulated sites in PP2A-B55 interactors to regulated sites in proteins not directly bound to PP2A-B55 (Fig. S7B).

Collectively, our integrated data establish a resource for understanding PP1 and PP2A-B55 dephosphorylation and provide evidence that the inherent dephosphorylation preferences of phosphatases is further shaped by binding to interactors.

### Regulation of PP1 RVxF interactions during mitotic exit

Our network analysis revealed a central role of RVxF motifs in mediating PP1 dephosphorylation activity and specificity. To determine if PP1 interaction with RVxF motifs is regulated during mitosis to influence dephosphorylation, we first obtained a more detailed quantitative characterization of the motif. The RVxF architecture is best described as Rx^0-1^VxF and we characterized this motif by synthesizing a library of peptides directly on amine-functionalized MRBLEs prepared using a simplified process, again with a 1:1 linkage between synthesized peptide identity and embedded spectral code [9, 17, 34]. After incubating bead-bound peptide libraries with PP1, an unlabeled anti-PP1 primary antibody, and an Alexa 647- labeled secondary antibody, we washed beads and imaged again in both lanthanide and fluorescence channels to identify peptides and quantify the amount of PP1 bound (Fig. 5A). Repeating these experiments over 8 different PP1 concentrations enabled acquisition of concentration-dependent binding curves; fitting the mean intensity of all beads with a given code at different concentrations of PP1γ and fitting to a Langmuir isotherm made it possible to quantify absolute affinities (Figs. 5B and S9). We synthesized two different sets of peptides derived from the Nipp1 peptide, either AKNSRVTFSEDDEII (RVxF) or AKNRAVTFSEDDEII (RxVxF) and systematically changed amino acids at specific positions in and around the motif (Fig 5B-D). In addition, we studied the effect of phosphorylation at different positions, as this has been shown to regulate PP1 binding (Fig. 5D, phosphosites indicated with red squares)[8, 35]. Measured affinities identified substitutions that both enhanced and ablated binding (Fig. 5B) and were highly reproducible across experimental replicates over three orders of magnitude (Fig. 5C).

**Figure 5.**
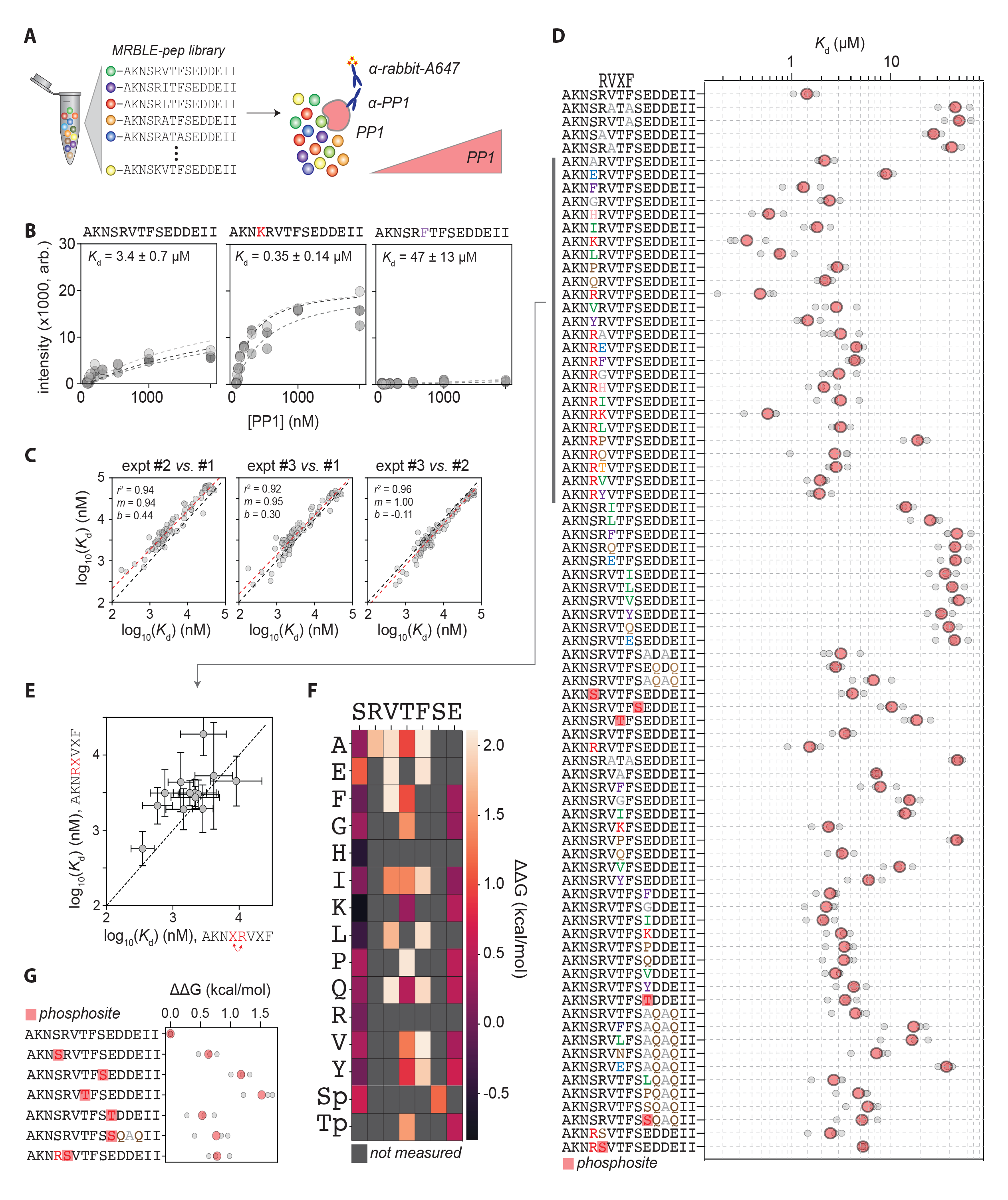
RVxF motif analysis. **A**) Overview of MRBLE-Pep experiments to quantify PP1 binding via incubation of bead-bound peptide libraries with different concentrations of PP1 pre-incubated with an anti-PP1 primary antibody and Alexa 647-labeled secondary antibody. **B**) Examples of MRBLE-Pep concentration-dependent binding measurements (markers) and Langmuir isotherm fits (dashed lines). Different marker and line shades indicated different experimental replicates; annotated *K*_d_s indicate the median and standard deviation of returned fit values across experimental replicates. **C**) Comparison of log-transformed fitted *K*_d_ values between experimental replicates. Black dashed line indicates the identity line, red dashed line indicates the best fit linear regression of log-transformed values, and annotation denotes linear regression results. **D**) MRBLE-pep affinities (*K*_d_s) for each peptide sequence. Gray markers indicate Kds returned from individual experiments; red markers indicate median values across experiments. The position of the core RVxF motif is indicated above the peptide sequences and red squares denote phosphorylated residues. **E**) Impacts of the same residue substitution at either the first x position within the RxVxF motif (*y* axis) or the −1 position within the RVxF motif (*x* axis). Black dashed line indicates identity line; markers denote mean and error bars denote standard deviation across experimental replicates. **F**) Heat maps showing impacts of different single amino acid substitutions at positions within and immediately surrounding the RVxF motif. Differences in binding affinities (ΔΔGs) are computed relative to the ‘AKNSRVTFSEDDEII’ reference sequence; grey boxes denote mutations not measured in this experiment. **G**) Impacts of phosphorylation at specific sites on binding affinities (again computed relative to the ‘AKNSRVTFSEDDEII’ reference sequence).

It has previously been established that the presence of an additional amino acid between the 0 and +1 position is tolerated and our data confirms that there are only small changes in binding energy between RVxF and RxVxF peptide variants (Fig. 5D-E). MRBLE-pep data confirm that single amino acid substitutions at the −1 position relative to RVxF have similar impacts as the same substitutions at the first x position within the RxVxF motif, although RxVxF peptides are consistently bound with slightly lower affinities (Figs. 5D, G). Most amino acids in the −1 position (in the context of RVxF) are tolerated with a notable preference for basic residues (K, R or H) (reduced ΔΔG), while a loss of binding occurs for acidic residues (E) (Fig. 5E); phosphorylation of the +1 position reduces binding and most substitutions away from the E at the +2 position lead to a small loss of binding (Fig. 5E). While the valine can be substituted by a leucine or isoleucine, the binding affinity is substantially weaker (Figs. 5D-E). Most substitutions at the first x position within the RxVxF motif have little impact on binding with the exception of a lysine residue; prolines inside the core RVxF or RxVxF are strongly disfavored (Fig. 5D-F). All phosphorylation sites strongly negatively regulated PP1 binding with increases in binding energy at positions from −1 to +2 and +4 (Fig 5F). We validated this by ITC using a number of model peptides (Fig. S10A-B).

Given the strong negative impact on PP1 binding to RVxF motifs by phosphorylation, we analysed our PP1 dephosphorylome for regulated sites in RVxF motifs. This revealed 15 RVxF motifs that were dephosphorylated by PP1 (Fig. S10C). Using several phosphopeptides, we confirmed that PP1 could indeed dephosphorylate RVxF motifs (Fig. S10D). The dephosphorylation of a Nipp1 peptide with a mutated RVxF motif (RVpTF to RApTA) was still efficiently dephosphorylated by PP1 suggesting that dephosphorylation of RVxF motifs is independent of PP1 binding to the motif (Fig. S10D).

Collectively, this suggests that PP1 can stimulate its own association with RVxF motifs at mitotic exit, potentially to amplify PP1 activity on specific substrates.

## Discussion

Our understanding of phosphatases in the control of cellular signaling pathways is still limited, partly due to technical limitations. Here we describe novel approaches to study phosphatases and use these to get new insight into mitotic exit regulation by PP1 and PP2A-B55. Our analysis integrates, for the first time, phosphatase dephosphorylation site motif preferences on a global scale on endogenous substrates with high-throughput *in vitro* dephosphorylation data.

We extended the use of our MRBLE assay from protein-peptide binding experiments to multiplexed peptide dephosphorylation assays. The ability to detect the phosphorylation status of peptides attached to encoded beads greatly increases the use of the MRBLE technology and should easily be adaptable to kinase assays. The small reaction volume allows the use of difficult to produce recombinant holoenzymes, and the bead technology offers an advantage compared to microarrays in terms of limited protein amounts needed and kinetic information. The systematic MRBLE:Dephos measurements allowed us for the first time to observe crosstalk between N-terminal and C-terminal residues surrounding the phosphorylation site. The effect of negative dephosphorylation determinants can be alleviated by positive determinants elsewhere in the peptide sequence, underscoring the complexity of dephosphorylation determinants.

Our analysis of PP1 and PP2A-B55 supports that protein phosphatases, like protein kinases [2], prefer specific phosphoacceptors and amino acid motifs around the phosphorylation site. We show the biological significance of this using INCENP dephosphorylation as an example. Our work expands on earlier observations on PP1 and PP2A complexes using a limited number of synthetic model peptides [21, 36-41]. We find that PP1 and PP2A-B55 preferentially dephosphorylate phospho-threonine over phospho-serine residues and that a Pro residue at +1 reduces dephosphorylation kinetics across a large panel of peptides, consistent with recent and earlier data [21, 28]. Interestingly, our integration of interactomes and regulated sites reveals that direct binding of a phosphatase allows it to dephosphorylate suboptimal sites such as SP sites close to the RVxF motif in KI67.

In addition, we identified preferences of PP1 and PP2A-B55 for specific amino acid motifs surrounding the phosphoacceptor with PP1 preferring basic residues at −1 and −2 positions while this is not a preference displayed by PP2A-B55. In contrast, PP2A-B55 prefers an acidic residue or glycine residue at −1 while acidic residues C-terminal to the phosphorylation site was deselected against. The difference in the stimulation of dephosphorylation by basic amino acids at −1 and −2 could stem from differences in the acidic grooves on the surface of the catalytic subunits. While conserved in PP1 and PP2A, the PP1 acidic groove is more extensive, promoting interactions with basophilic phosphorylation site motifs [42, 43].

Dephosphorylation motif preferences of PPPs have implications for understanding disease mutations around phosphorylation sites, which might affect not only phosphorylation but also dephosphorylation. Future efforts in establishing dephosphorylation signatures for all phosphatases will be important and complement the current knowledge we have of kinase consensus sites [2], allowing better interpretation of phosphorylation site data. However, ultimately understanding of phospho-dependent signaling will require a full understanding of phosphorylation/dephosphorylation motif signatures and docking sites for kinases and phosphatases. The assays described here will enable such discoveries on a global scale.

## Methods and materials

### MRBLE Production and Peptide Synthesis

We generated PEG-DA Hydrogel beads with 96 different lanthanide-nanoparticle (LNP) based codes as described previously. We premixed a solution to generate droplets containing ratios of LNP to generate a code. 21 % PEG-DA 575, 3% LAP (39.2 mg/ml) and Pent-4- enylamine were mixed with the appropriate amount (Table S1) of Eu (5 % v/v YVO_4_:Ln (50mg/mL), Dy, Sm, or Tm master mix (16.3 % v/v YVO_4_:Ln (50mg/mL). Pent-4-enylamine (0.09% v/v) was added to HFE7500 oil supplemented with custom-made ionic Krytox (2.2 wt%). A microfluidic PDMS parallel flow focusser with two syringe pumps was used to generated water-in-oil emulsions that were collected in wells containing HFE7500 oil The polymerization of the PEG-DA was induced with UV-light ((IntelliRay, UV0338) for 2 min at 100% amplitude (7” away from the lamp, power = ∼50–60 mW/cm^2^). The beads were then washed with 2 mL DMF, 2 mL DCM, 2 mL MeOH, 2 mL H_2_O and 2 mL PBS-T. Prior to peptide synthesis, ∼10000 beads were washed with 3 mL MeOH, 3 mL DCM and 3 mL DMF. The beads were sonicated for 20 minutes in a water bath sonicater and mixed overnight. Prior to HT-SPPS, the first amino acid was attached to the amino group on the beads. To each vial containing beads (∼1 mg, 0.32 mmol/g loading capacity) with a given code, 400 uL DMF with 5 eq Fmoc-Glycine, 5 eq DIC and 5 eq DIPEA was added. The reaction was mixed overnight and repeated. After extensive washing with DMF, MeOH, and DMF, free amine groups were capped with a mixture of Acetic-Anhydride and 25% Pyridine for 15 minutes. Fmoc glycine was deprotected using 20% 4-methylpiperidine before peptides were synthesized on a HT-SPPS (Microsyntech Syro II) using double coupling for 20 minutes and 10 eq amino acid, 9.8 eq HCTU and 20 eq NMM. At difficult coupling steps (e.g. for phosphorylated serine or threonine), an additional capping step was introduced. The peptides were Fmoc deprotected using 4-methylpiperidin and sidechain deprotected using freshly prepared Reagent B (95 % TFA, 2.5% TIPS and 2.5% H_2_O.

### MRBLE:Dephos

We synthesized 95 peptides (15 amino acids) on encoded beads as described above (Table S2). The beads from each code were mixed and incubated with PMP Buffer (50 mM HEPES, 100 mM NaCl, 2 mM DTT, 0.01% Brij 35, pH 7.5) supplemented with 1 mM MnCl_2_ and 0.2% BSA for 10 minutes at 30°C and 800 rpm. The mixed beads were divided into tubes and PMP Buffer with 1 mM MnCl was added along with either PP1 (600 ng), PP2A-B55 (5 ul of elution) or Lambda phosphatase (2 ul). The reaction was stopped by adding 1 ul Calyculin A (Cell Signaling). The buffer was removed, 100 μl TBS-T supplemented with Calyculin A was added, and the beads were then transferred to a 1.5 mL Eppendorf tube. The beads were sonicated for 10 minutes in a water bath sonicator, the TBS-T was exchanged with 500 μl pIMAGO blocking buffer, and beads were incubated for 30 minutes at RT at 1450 rpm with a 15 sec. on/off interval. The beads were washed once with pIMAGO Buffer, and 500 μl fresh pIMAGO Buffer containing 20 ul pIMAGO was added and incubated for 1 hour at RT at 1450 rpm 15 sec on/off. Wash beads three times with 500 uL washing buffer (2 min., 5 min., 2min.) at 1450 rpm with a 15 sec. on/off and one time with TBS-T. Beads were then incubated with DyLight 650- labelled Streptavidin for 30 min. at 1450 rpm with a 15 sec. on/off interval. The beads were washed three times with TBS-T prior to imaging as described previously [44]. MRBLE:Dephos experiments measured per-bead Cy5 intensities at the following timepoints: 0 minutes (initial intensity prior to dephosphorylation), 15 minutes, 30 minutes, 60 minutes, 120 minutes, and 240 minutes.

### MRBLE:Dephos Analysis

To analyze MRBLE:Dephos data, we first computed the mean Cy5 intensity for all beads of a given code/peptide sequence at each timepoint. To quantify the degree to which each peptide was dephosphorylated in a given experiment, we then: (1) computed a per-code final value by averaging over the last 3 timepoints (over which signals were typically relatively constant), (2) computed the fraction of phosphate groups remaining by dividing this final value by the mean Cy5 intensity of each code prior to adding phosphatases (at timepoint 0) (‘ratio’), and then (3) subtracting this ratio from 1 such that larger scores indicated more phosphorylation (‘dephosphorylation score’). To report a final per-peptide dephosphorylation score, we computed the mean and standard deviation of dephosphorylation scores across 3 independent replicates. All raw data for all experiments and code used to process these data are available in an OSF repository associated with the paper (https://osf.io/gys5d/).

### MRBLE:Pep

We synthesized 70 peptides (15 amino acids) on encoded Beads as described above (Table S3). PP1 concentration-dependent binding to peptides on MRBLEs was done as described previously (Hein et al.). In short, ∼700 beads per code were mixed in and blocked in 700 uL PBS-T with 5% BSA. PP1 (2uM) was preincubated in binding buffer (50 mM Tris pH = 7.5, 150 mM NaCl, 0.1% TWEEN 20) with 1 uM primary (a-PPP1CC rabbit polyclonal, Bethyl A300-906A) and 1 uM goat anti-rabbit IgG Alexa Fluor 647 (Thermo Fischer, A-21245). The beads were divided into 7 tubes and the appropriate concentration of PP1 antibody mixture was added at a final volume of 100 ul. After 24 hours incubation time at 4 C the buffer was exchanged once and the beads were resuspended in 20 ul PBS-T prior to imaging. Bead imaging has been performed as described previously. Each MRBLE:Pep experiment quantified Cy5 intensities for PP1 incubated with beads at 0, 15, 31, 62, 125, 250, 500, 1000, and 2000 nM concentrations.In the first experiment the two sets of beads (codes 1-48 and 49 to 96) were done separately with above described conditions. For the second and third experiment the two sets were pooled and the volume was increased to 200 μl with the same concentration as in the first experiment.

### MRBLE:Pep Analysis

To begin MRBLE:Pep analysis, we first computed the median Cy5 intensity for all beads with a given code/peptide sequence for each PP1 concentration. To allow global fitting of Langmuir isotherms to determine *K*_d_ values even for peptides that do not saturate, we assume that all peptides would bind PP1 at the same stoichiometry such that they would all saturate at the same final value. To quantify this value, we identified all peptides that exhibited saturation at high PP1 concentrations (*i.e.* had final Cy5 intensities > 12,000) and locally fit concentration-dependent binding behavior to the Langmuir isotherm equation:

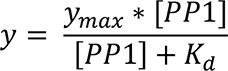

We then computed the mean *y*_max_ value across all of these peptides and refit concentration-dependent binding curves for all peptide sequences to the Langmuir isotherm equation using this experimentally determined *y*_max_ value for all peptide sequences and fitting only the *K*_d_ term. To determine a final reported *K*_d_, we then averaged these fitted *K*_d_ values across all 3 experiments. All raw data and Jupyter notebooks for the analysis are provided in the OSF repository associated with the paper (https://osf.io/gys5d/).

### Phosphorylation of GST-Arpp19 with MASTL and INCENP with Cdk1

Thiophosphorylated GST-Arpp19 was generated as described previously [28]. In short, purified GST, GST-Arpp19^wt^ or GST-Arpp19^S62A^ was incubated with purified 3xFLAG-MASTL kinase in protein kinase buffer (50 mM Tris-Hcl pH 7.5, 10 mM MgCl2, 0.1 mM EDTA, 2 mM DTT, 0.01% Brij 35) with 500 μM ATP and 1 μCi [γ-32P] ATP at 30°C for 30 min, or with 500 μM ATPγS at 30°C for 60 min for pull-down experiments.

Kinase reactions were carried out in non-stick 1.5 mL tubes. 15 μg of GST-INCENP wt or S61E were incubated with the indicated amounts of CDK1-CyclinB1 (Sigma #SRP5009) in 50 μL reactions in kinase buffer (50 mM Tris– HCl pH 7.5, 10 mM MgCl2, 0.1 mM EDTA, 2 mM DTT, 0.01% Brij 35) with 500 μM ATP and 1 μCi (γ-32P)-ATP (Perkin Elmer) at 30°C for 60 min. Reactions were stopped by the addition of 10 μM RO-3306 (Calbiochem). Samples of ∼ 2μg were were separated by SDS–PAGE. Gels were dried, exposed for 3 days, and imaged on Typhoon FL 950 (GE Healthcare). Analyses and quantifications were carried out in ImageJ.

### Lysate Assay

Phosphatase Inhibition in cell lysates was performed as described previously. Double Thymidine synchronized cells were arrested in Nocodazole for 14 hours. Cells were collected by mitotic shake off and washed with PBS. Room temperature lysis Buffer (150 mmol NaCl, 25 mmol Tris, 0.1% NP40, complete protease Inhibitor) with 50 ug of GST-tagged thiophosphorylated Arppt19 or Nipp1 central domain was used to resuspend cell pellets and immediately transferred to a thermomixer at 600 rpm for 5 minutes. The reaction was stopped by adding the same amount of lysis buffer with 2x concentration of PhosStop inhibitor tablets. The lysate was cleared for 15minutes at 20000g at 4 C and the supernatant was snap frozen.

### Protein Purification

His-PP1γ, GST-Arpp19, GST-INCENP and His-Nipp1 was expressed in BL21(DE3) overnight at 18 degrees. Cells where lysed and following lysis the supernatant was clarified by centrifugation and loaded on an affinity column. Following washing the samples where eluted and fractions analysed by SDS-PAGE. Peak fractions where pooled and concentrated and further purified using size exclusion chromatography. Proteins where validated by mass spectrometry.

PP2A-B55α holoenzymes was purified from HeLa cells stably expressing FLAG tagged B55α as previously described[12].

### RNAi and plasmid tranfection

RNAi Max (Life Technologies) was used to deplete endogenous proteins. INCENP (5’- AGUCCUUUAUUAAGCGCAA-3’) silencer select siRNA was obtained from Thermo Fischer). For each well in a 6 Well dishes 250 ul Opti-MEM with 3 ul RNAiMax and 250 ul Opti-MEM with 2.5 ul siRNA were mixed after 5 minutes and after 20 minutes added to 1 ml Opti-MEM. The reverse transfection into IBIDI dishes was performed as follows: 10% of the cells from each 6 Well dish were resuspended in 200 ul Opti-MEM and a transfection mix containing 1 ul siRNA Max and 0.25 ul siRNA in 50 ul Opti-Mem was added. 200 ng pcDNA5 FRT/TO YFP or YFP-INCNEP WT or YFP-INCENP S61E was added to 200 ul jetOptimus Buffer mixed and supplemented with 1 ul jetOptimus transfection reagent. After 10 minutes the tranfection mix was added to 1.8 ml DMEM with 10% FBS.

### Live cell

HeLa cells were seeded in a 6 Well cell culture dish in complete DMEM and synchronized using a double thymidine block. Before the first addition of thymidine cells were transfected with plasmids using jetOptimus and subsequently with siRNA Max. The transfection mix was supplemented with DMEM, FBS and Thymidine. The following day, after release from thymidine cells were reverse transfected with siRNA using siRNAMax and seeded into 8Well Ibidi microscopy slides. 8 hours after release the transfection mix was supplemented with DMEM, FBS (final concentration 10%) and thymidine. The following day the cells were released from thymidine into complete DMEM media. After 4 hours the medium was changed to L15 supplemented with 10% FBS and the slide was mounted on a Delta Vision Elite microscope. The cells were imaged for 18 to 22 hours in a 8 minute interval using a 40x, 1.35 NA, working distance 0.10 objective at 37C. The data was analyzed using softWoRx and Image J. Statistical analysis was performed using the Mann-Whitney test in Prism.

### Quantitative Western Blot

Cell lysates were analyzed using infrared Western blot technology (Li-Cor Bioscience). After SDS-PAGE separation and western blot onto Immobilion FL membrane, proteins were detected using indicated primary antibody and IRDye 800 or 680 secondary antibodies. The Odyssey CLx imaging system was used to scan the membranes.

### Phosphate Release Assay

In vitro phosphatase assays were performed with E.coli expressed PP1γ or PP2A-B55α purified from HeLa cell extracts as described previous. Phosphorylated peptides (Peptide 2.0) at a concenentrations 120 uM were incubated with 80 ng PP1 or PP2A-B55 for 20 minutes at 30 C in phosphatase Buffer (50 mM Tris pH 7.4, 1 mM MnCl_2_, 150 mM NaCl). The PiColorLock inorganic phosohate detection kit (abcam) was used to detect the amount of phosphate released from the phosphorylated peptides.

### Proximity dependent ligation and affinity purification of phosphatases

PP1 and B55 cDNAs where cloned into modified versions of pcDNA5/FRT/TO (Invitrogen) to make N terminal fusions to YFP, BioID or TurboID. Stable cell lines were generated into HeLa/FRT/TRex cells (kind gift from S. Taylor) and for YFP affinity purification cells were synchronized with Thymidine and released into nocodazole containing media to arrest them in mitosis. Cells were resuspended in lysis buffer (50 mM NaCl, 50 mM Tris 7,4, 1 mM EDTA, 0,1% NP40, 1mM DTT, protease inhibitor cocktail) and following sonication the lysate was clarified by centrifugation. Cell lysate was incubated with GFP-Trap beads (Chromotek) for 1 hour and washed 3 x 1ml with lysis buffer and resuspended in SDS-PAGE loading buffer.

For proximity dependent ligation using BioID cells were synchronized as above, and biotin added overnight to the medium and biotinylated proteins purified as described in [45]. As control we used samples not treated with biotin. For TurboID analysis cells were synchronized in S-phase using thymidine and aphidicolin and cells released from this block by washing with fresh media. 5 hours after release biotin was added for 1 hour and cells harvested and processed as for BioID samples.

### Quantitative TMT phosphoproteomics analysis

Thiophosphoryalted GST-Arpp19^wt^ or GST-Arpp19^S62A^ and NIPP or NIPPmut treated cell lysates were snap-frozen and proteins were acetone precipitated [12]. The precipitated proteins were digested overnight with trypsin (1:100 w/w) at 37°C. Digests were desalted using C_18_ solid-phase extraction cartridges (ThermoFischer Scientific) and dried by vacuum centrifugation. Phospho-peptides were enriched in these samples using High-Select™ Fe-NTA Phosphopeptide Enrichment Kit according to manufacturer’s protocol (ThermoFisher Scientific). Phosphopeptides were resuspended in 133 mM HEPES (SIGMA) pH 8.5 and TMT reagent (ThermoFisher Scientific) stored in dry acetonitrile (ACN) (Burdick & Jackson) was added followed by vortexing to mix reagent and peptides. After 1 hr at room temperature, an aliquot was withdrawn to check for labeling efficiency while the remaining reaction was stored at −80°C. Once labeling efficiency was confirmed to be at least 95%, each reaction was quenched with ammonium bicarbonate for 10 minutes, mixed, acidified with 20% TFA, and desalted. The desalted multiplex was dried by vacuum centrifugation and separated by offline pentafluorophenyl (PFP)-based reversed phase HPLC fractionation as previously described [46]. TMT-labeled samples were analyzed on an Orbitrap Fusion [47] mass spectrometer (ThermoScientific) equipped with an Easy-nLC 1000 (ThermoScientific). Peptides were resuspended in 8% methanol / 1% formic acid across a column (45 cm length, 100 μm inner diameter, ReproSil, C_18_ AQ 1.8 μm 120 Å pore) pulled in-house across a 2 hr gradient from 3% acetonitrile/0.0625% formic acid to 37% acetonitrile/0.0625% formic acid. The Orbitrap Fusion was operated in data-dependent, SPS-MS3 quantification mode [48, 49] wherein an Orbitrap MS1 scan was taken (scan range = 350 – 1200 m/z, R = 120K, AGC target = 3e5, max ion injection time = 100ms). Followed by data-dependent Orbitrap trap MS2 scans on the most abundant precursors for 3 seconds. Ion selection; charge state = 2: minimum intensity 2e5, precursor selection range 650-1200 m/z; charge state 3: minimum intensity 3e5, precursor selection range 525-1200 m/z; charge state 4 and 5: minimum intensity 5e5). Quadrupole isolation = 0.7 m/z, R = 30K, AGC target = 5e4, max ion injection time = 80ms, CID collision energy = 32%). Orbitrap MS3 scans for quantification (R = 50K, AGC target = 5e4, max ion injection time = 100ms, HCD collision energy = 65%, scan range = 110 – 750 m/z, synchronous precursors selected = 5).

The raw data files were searched using COMET (release version 2014.01) [50] against a target-decoy (reversed) [51] version of the human proteome sequence database (UniProt; downloaded 2/2020, 40704 entries of forward and reverse protein sequences) with a static mass of 229.162932 on peptide N-termini and lysines and 57.02146 Da on cysteines, and a variable mass of 15.99491 Da on methionines and 79.96633 Da on serines, threonines and tyrosines and filtered to a <1% FDR at the peptide level. Quantification of LC-MS/MS spectra was performed using in house developed software [12]. Phosphopeptide intensities were adjusted based on total TMT reporter ion intensity in each channel and log_2_ transformed. P-values were calculated using a two-tailed Student’s t-test assuming unequal variance.

### Label-free LC-MS/MS analysis

Pull-downs or proximity labelled samples were analyzed on a Q-Exactive Plus quadrupole Orbitrap mass spectrometer (ThermoScientific) equipped with an Easy-nLC 1000 (ThermoScientific) and nanospray source (ThermoScientific). Peptides were resuspended in 5% methanol / 1% formic acid and loaded on to a trap column (1 cm length, 100 μm inner diameter, ReproSil, C_18_ AQ 5 μm 120 Å pore (Dr. Maisch, Ammerbuch, Germany) vented to waste via a micro-tee and eluted across a fritless analytical resolving column (35 cm length, 100 μm inner diameter, ReproSil, C_18_ AQ 3 μm 120 Å pore) pulled in-house (Sutter P-2000, Sutter Instruments, San Francisco, CA) with a 45 minute gradient of 5-30% LC-MS buffer B (LC-MS buffer A: 0.0625% formic acid, 3% ACN; LC-MS buffer B: 0.0625% formic acid, 95% ACN).

The Q-Exactive Plus was set to perform an Orbitrap MS1 scan (R=70K; AGC target = 1e6) from 350 – 1500 m/z, followed by HCD MS2 spectra on the 10 most abundant precursor ions detected by Orbitrap scanning (R=17.5K; AGC target = 1e5; max ion time = 50ms) before repeating the cycle. Precursor ions were isolated for HCD by quadrupole isolation at width = 1 m/z and HCD fragmentation at 26 normalized collision energy (NCE). Charge state 2, 3, and 4 ions were selected for MS2. Precursor ions were added to a dynamic exclusion list +/- 20 ppm for 15 seconds. Raw data were searched using COMET (release version 2014.01) in high resolution mode [50] against a target-decoy (reversed) [51] version of the human proteome sequence database (UniProt; downloaded 2/2020, 40704 entries of forward and reverse protein sequences) with a precursor mass tolerance of +/- 1 Da and a fragment ion mass tolerance of 0.02 Da, and requiring fully tryptic peptides (K, R; not preceding P) with up to three mis-cleavages. Static modifications included carbamidomethylcysteine and variable modifications included: oxidized methionine. Searches were filtered using orthogonal measures including mass measurement accuracy (+/- 3 ppm), Xcorr for charges from +2 through +4, and dCn targeting a <1% FDR at the peptide level. Quantification of LC-MS/MS spectra was performed using MassChroQ [52] and the iBAQ method [53]. Missing values were imputed from a normal distribution in Perseus to enable statistical analysis and visualization by volcano plot [54]. Statistical analysis was carried out in Perseus by two-tailed Student’s t-test.

### IceLogo and WebLogo

The IceLogo and WebLogo were generated using the centered sequences of singly phosphorylated sites that were significantly regulated (p-value < 0.05; log_2_ ratio > 0.58 (1.5- fold)) by PP1 and B55 (Supp Table 1). WebLogo web server: www.weblogo.threeplusone.com [55]. IceLogo software is downloadable at www.github.com/compomics/icelogo [56].

### Antibodies

The following antibodies where used: GAPDH-rabbit-1:1000 (Santa Cruz; sc-25778), H3S10p – rabbit-1:1000 (Milipore #06-570), pan T(phos)P –mouse - 1:500 (Cell signaling #9391)

### ITC

Peptides were purchased from Peptide 2.0 Inc (Chantilly. VA, USA). The purity obtained in the synthesis was 95 – 98 % as determined by high performance liquid chromatography (HPLC) and subsequent analysis by mass spectrometry. Prior to ITC experiments both the protein and the peptides were extensively dialyzed against 50 mM sodium phosphate, 150 mM NaCl, 0.5 mM TCEP, pH 7.5. All ITC experiments were performed on an Auto-iTC200 instrument (Microcal, Malvern Instruments Ltd.) at 25 °C. Both peptide and PP1γ concentrations were determined using a spectrometer by measuring the absorbance at 280 nm and applying values for the extinction coefficients computed from the corresponding sequences by the ProtParam program (http://web.expasy.org/protparam/). The peptides at approximately 450 μM concentration were loaded into the syringe and titrated into the calorimetric cell containing the PP1γ at ∼ 35 μM. The reference cell was filled with distilled water. In all assays, the titration sequence consisted of a single 0.4 μl injection followed by 19 injections, 2 μl each, with 150 s spacing between injections to ensure that the thermal power returns to the baseline before the next injection. The stirring speed was 750 rpm. Control experiments with the peptides injected in the sample cell filled with buffer were carried out under the same experimental conditions. These control experiments showed heats of dilution negligible in all cases. The heats per injection normalized per mole of injectant *versus* the molar ratio [peptide]/[PP1γ] were fitted to a single-site model. Data were analysed with MicroCal PEAQ-ITC (version 1.1.0.1262) analysis software (Malvern Instruments Ltd.).

## Acknowledgements

Work at the Novo Nordisk Foundation Center for Protein Research is supported by grant NNF14CC0001 and JN is supported by grants from the Danish Cancer Society (R167-A10951- 17-S2), Independent Research Fund Denmark (DFF-4183-00388 and 8021-00101B) and Novo Nordisk Foundation (NNF16OC0022394 and NNF18OC0053124). JBH was funded by grant NNF17OC0025404 from the Novo Nordisk Foundation and the Stanford Bio-X Program. A.N.K was supported by grants from NIH/NIGMS (R35GM119455). PMF and JBH were supported by NIH grant DP2 GM123641 and PMF is a Chan Zuckerberg Biohub Investigator. We thank the protein production facility at NNF CPR for support with the project and the Cyert lab for discussions.

## Author contributions

JBH conducted all MRBLE experiments, lysate inhibition assays and INCENP experiments. HTN, IN and AK performed mass spectrometry analysis and analysis of this data. DHG prepared proteins for lysate experiments and helped with these experiments. YF helped with the initial production of MRBLE beads. BLM performed ITC measurements. PMF performed analysis of MRBLE data and ND helped with analysis of RVxF motifs. JBH, JN, AK and PMF drafted manuscript.

## Supplementary materials

SI word file: Contains 10 supplemental figures with legends

Table 1: Regulated phosphorylation sites from mitotic lysate screens

Table 2: Interactomes for PP1 and PP2A-B55

