## Supplemental Information for "Global substrate identification and high throughput *in vitro* dephosphorylation reactions uncover PP1 and PP2A-B55 specificity principles"

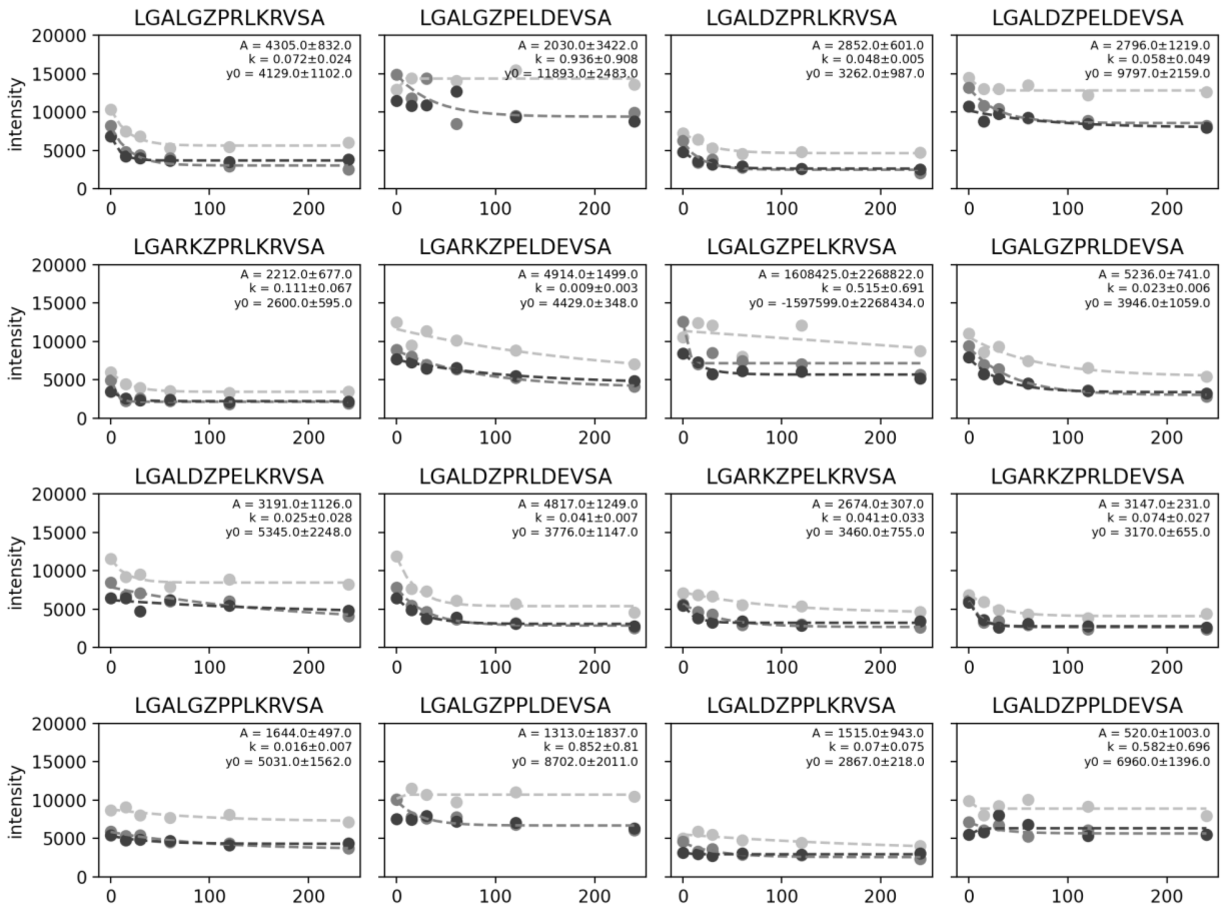


**Fig. S1**. Measured DyLight 650 intensities over time for MRBLE-bound peptides after incubation with PP1. Each marker shade denotes a different experimental replicate; dotted lines indicate a single exponential fit within each experimental replicate. Annotations report mean exponential fit parameters ± standard deviation across 3 independent replicates.

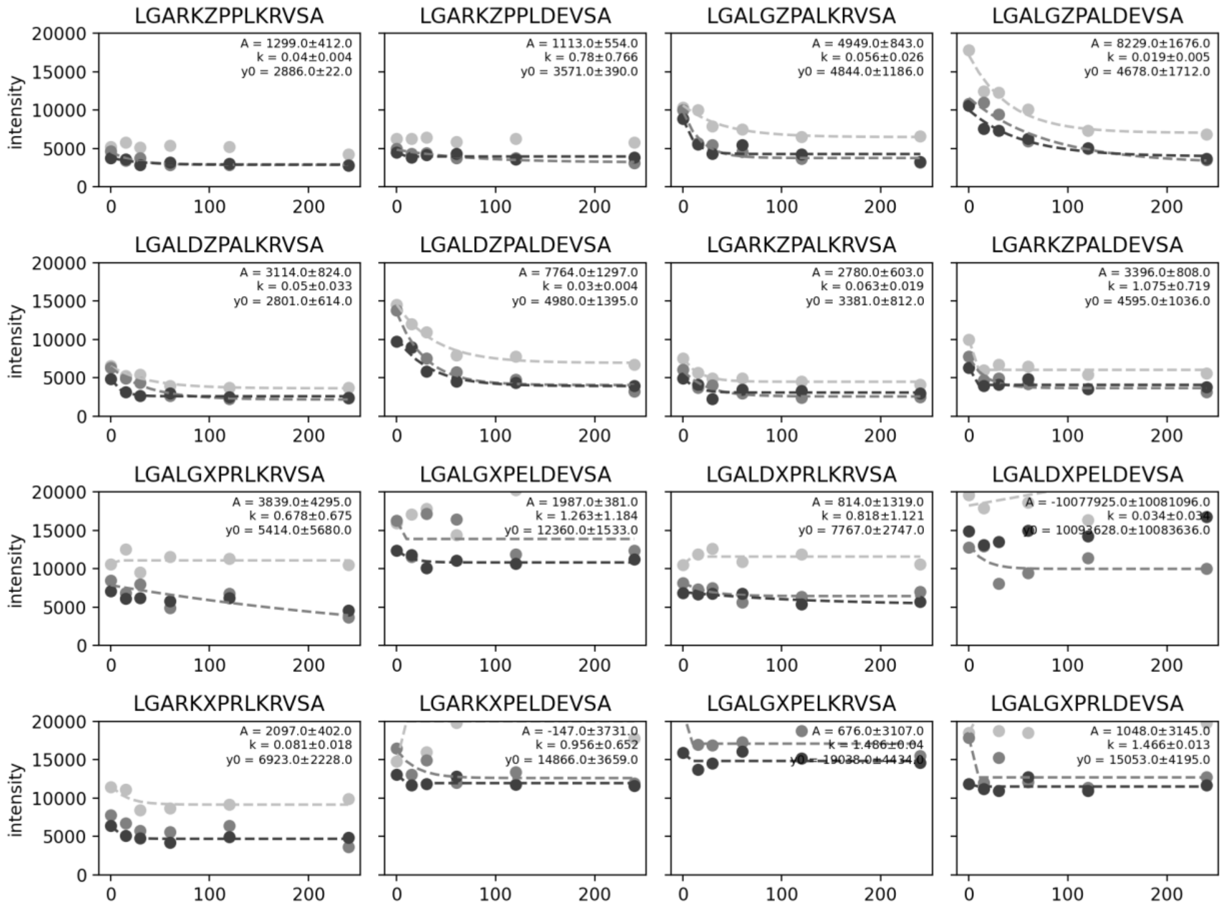


**Fig. S1 (continued).**

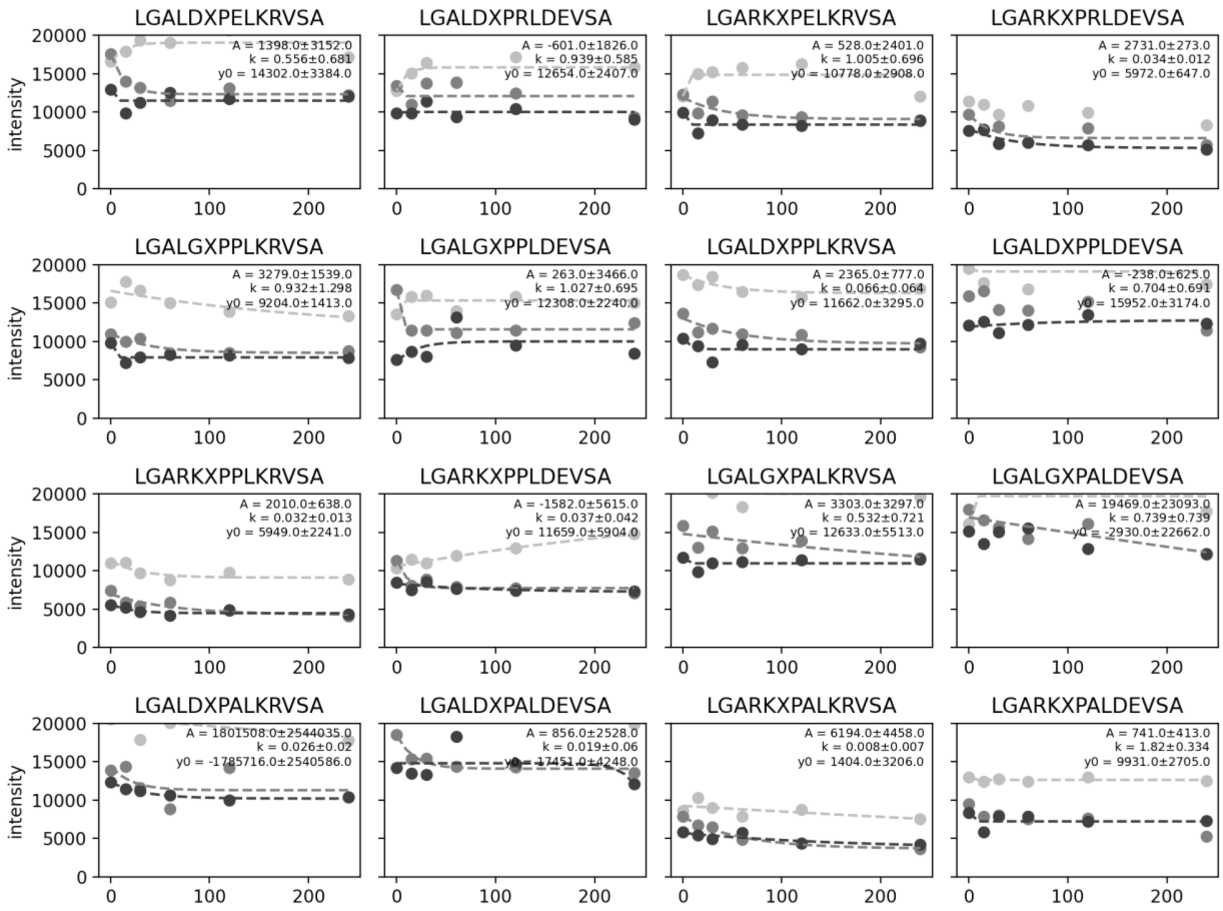


**Fig. S1 (continued).**

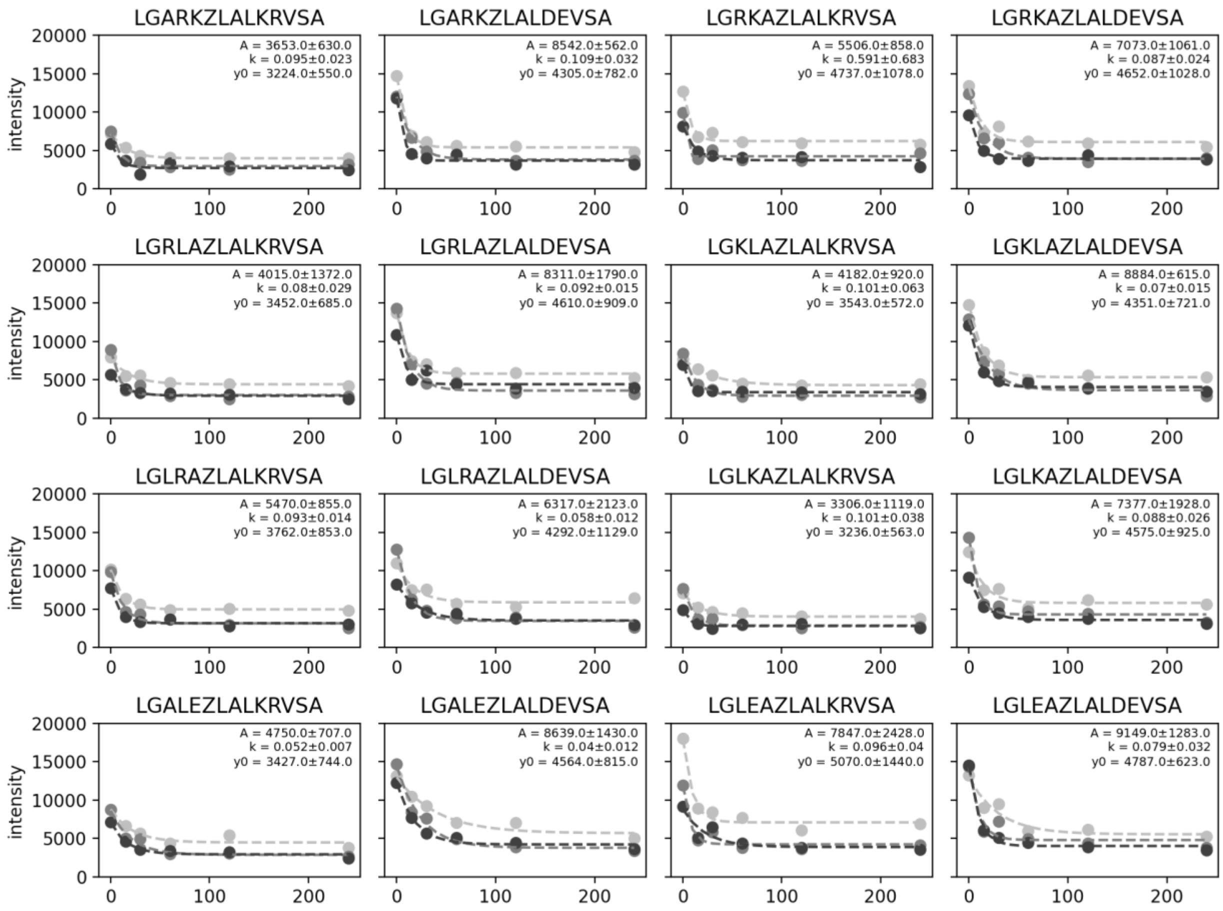


**Fig. S1 (continued).**

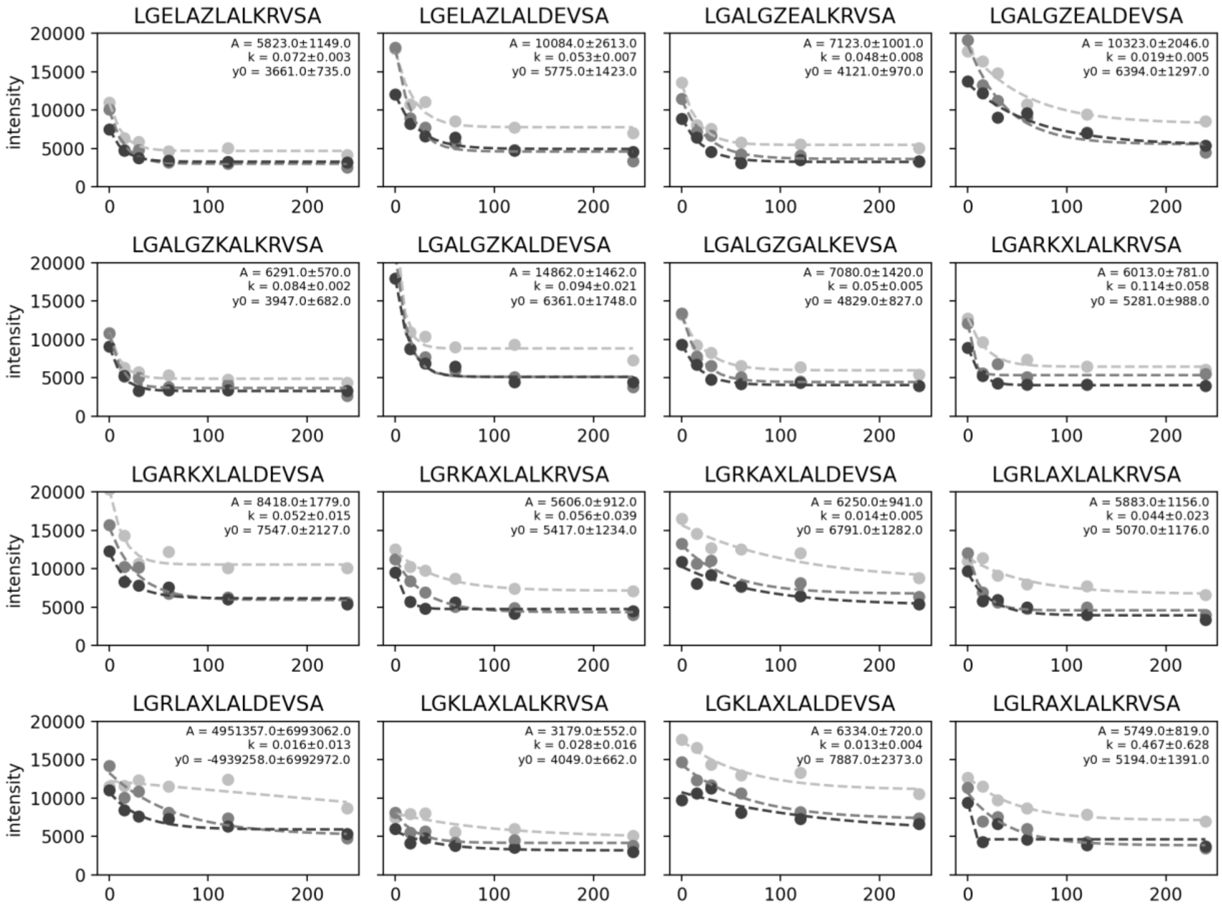


**Fig. S1 (continued).**

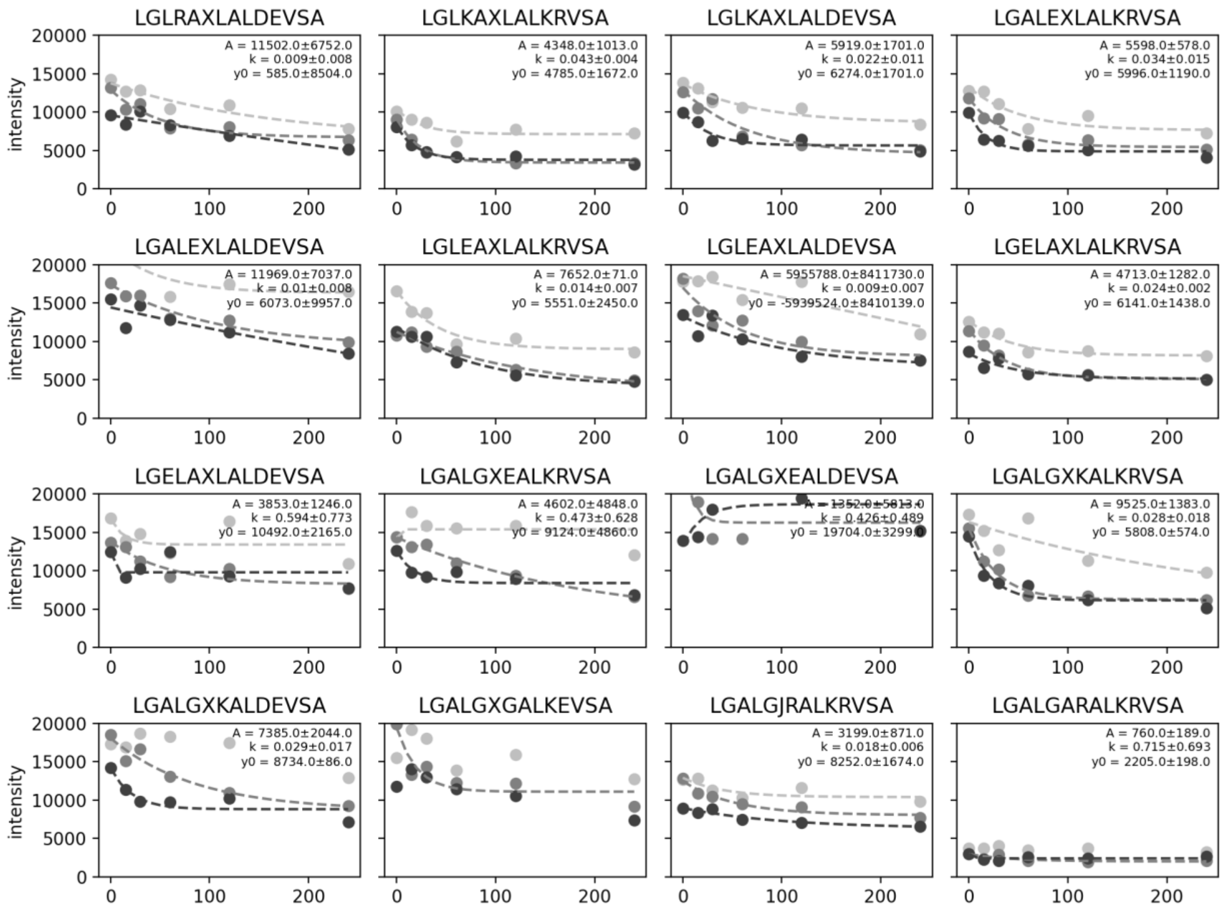


**Fig. S1 (continued).**

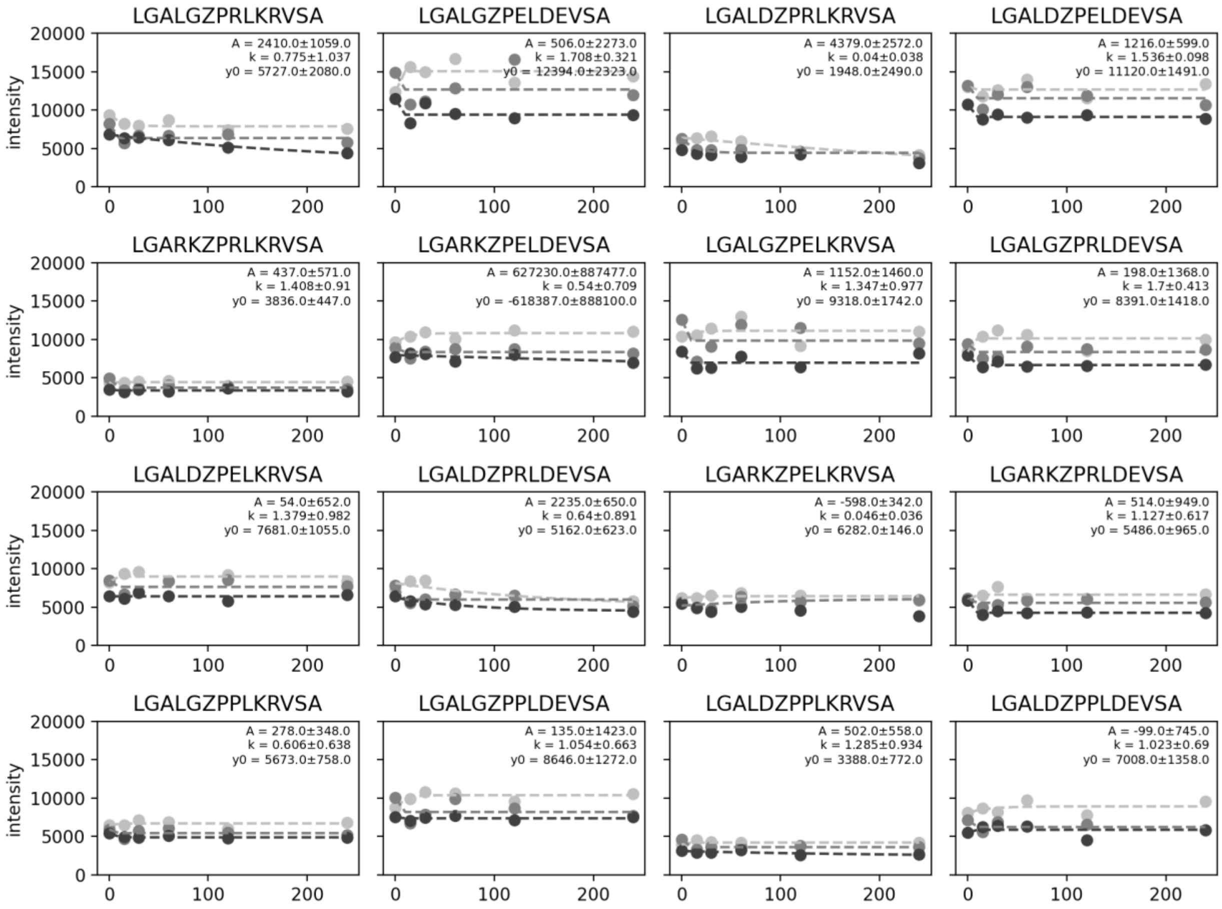


**Fig. S2.** Measured DyLight 650 intensities over time for MRBLE-bound peptides after incubation with PP2A-B55. Each marker shade denotes a different experimental replicate; dotted lines indicate a single exponential fit within each experimental replicate. Annotations report mean exponential fit parameters across 3 independent replicates.

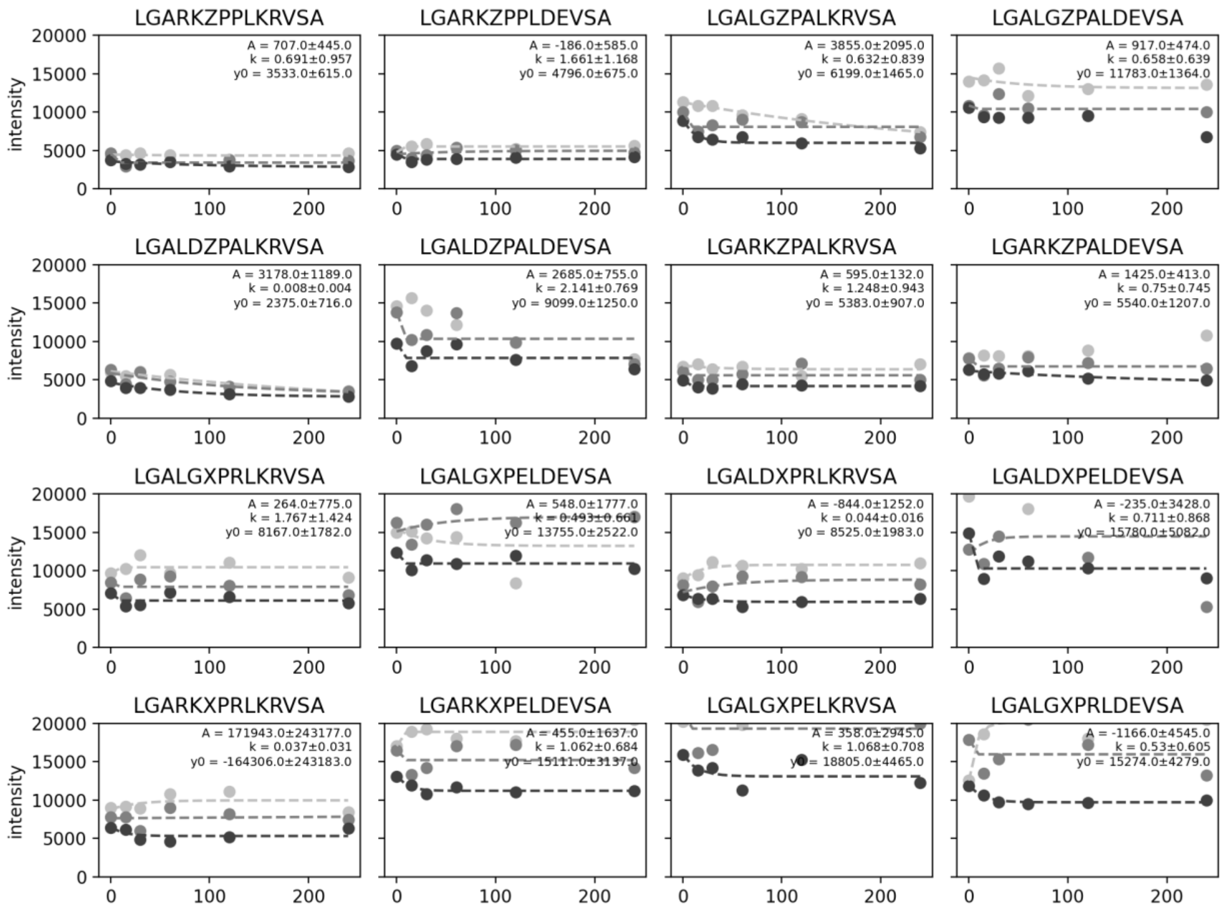

**Fig. S2 (continued).**

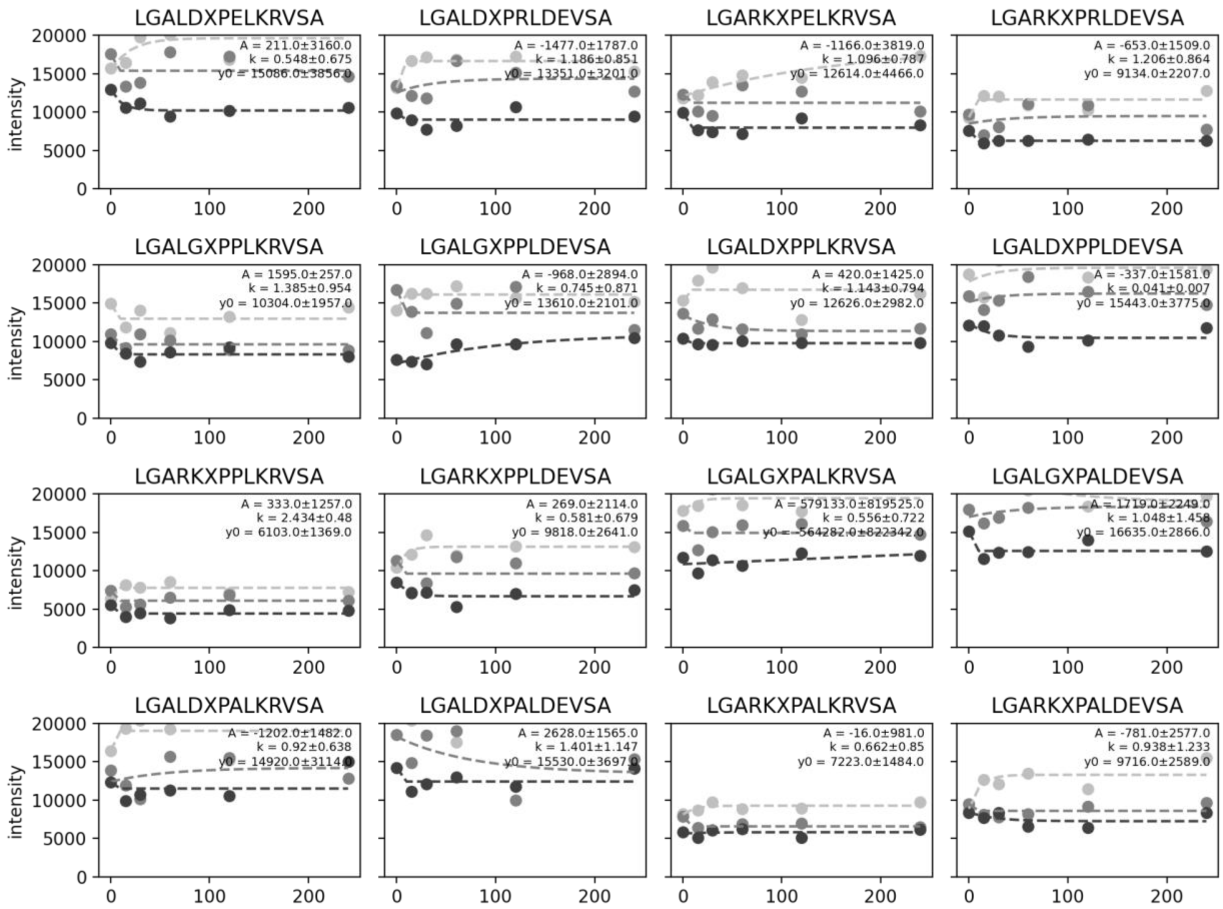


**Fig. S2 (continued).**

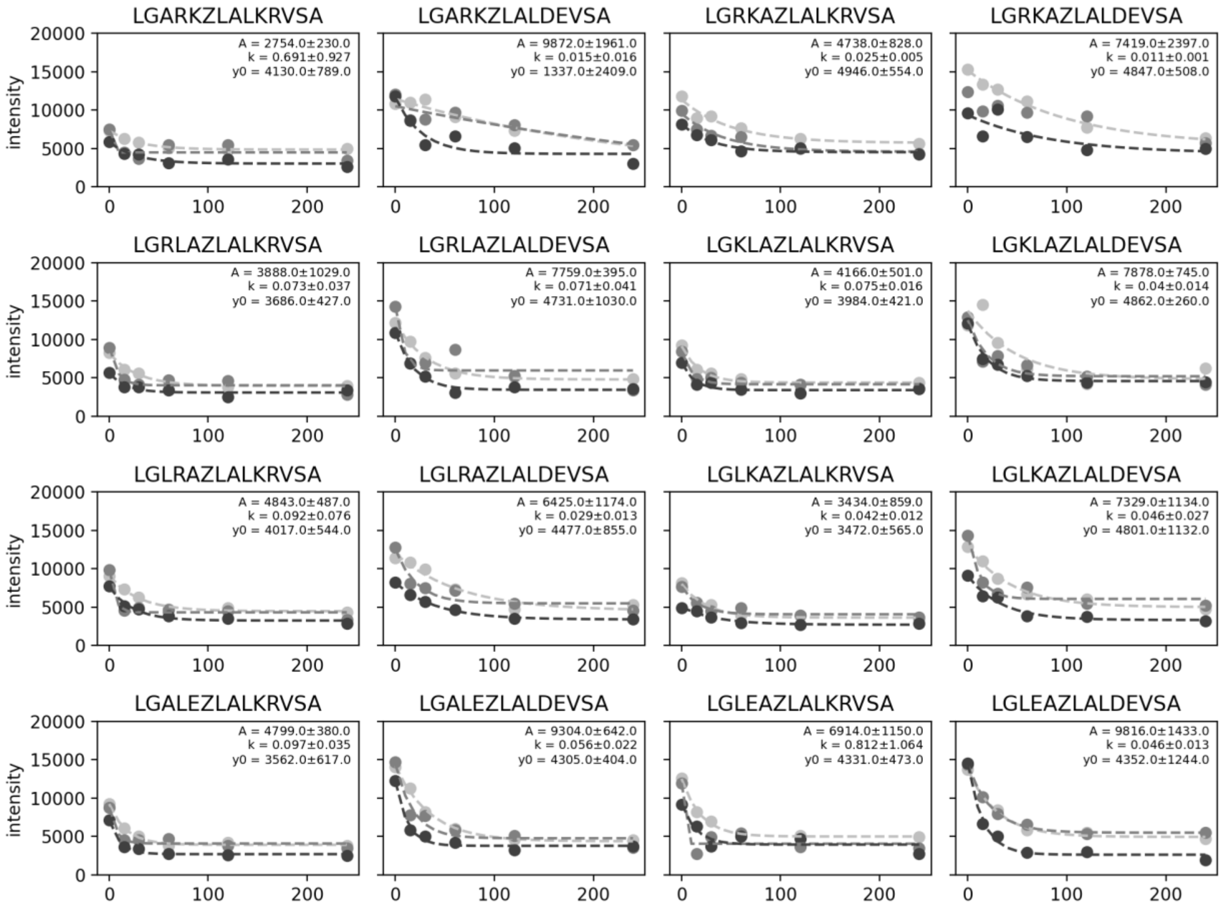


**Fig. S2 (continued).**

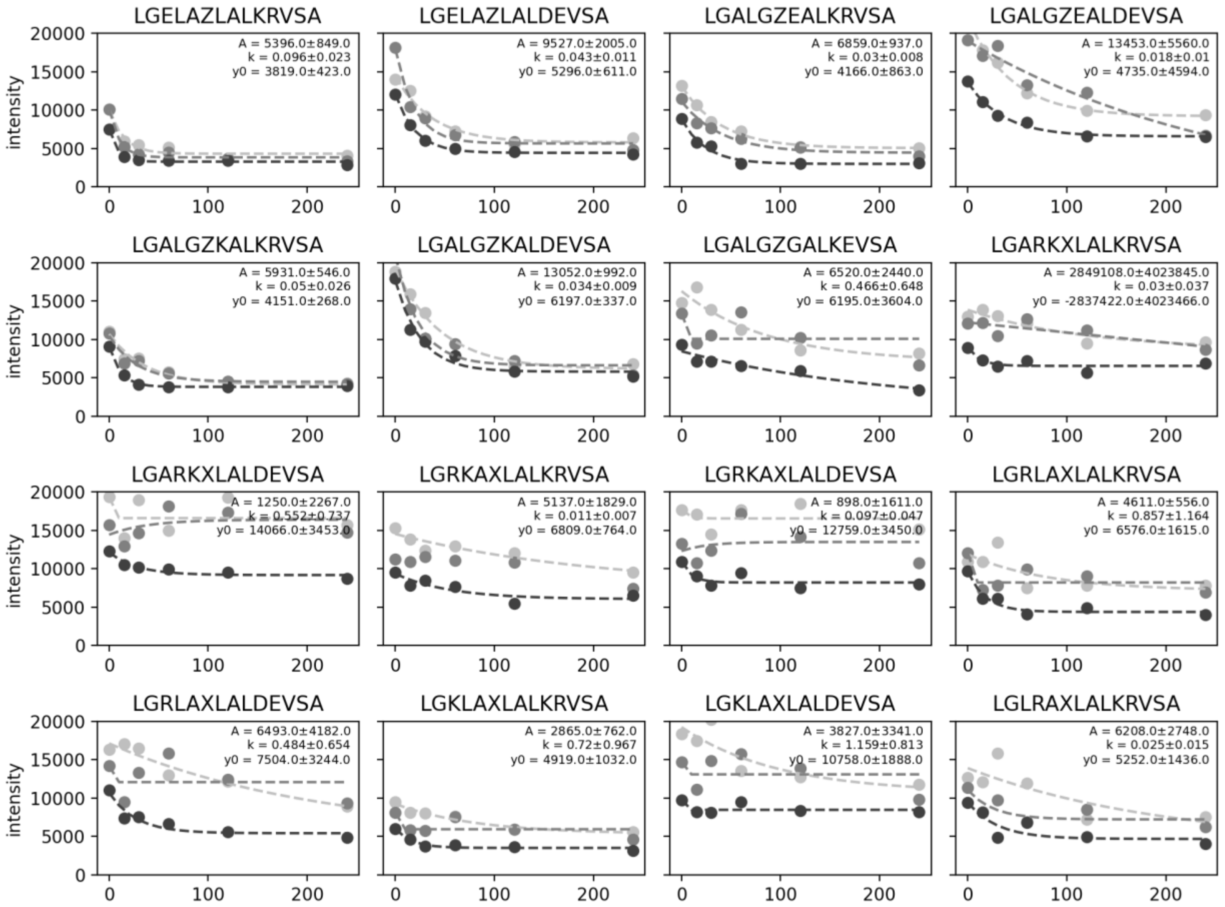


**Fig. S2 (continued).**

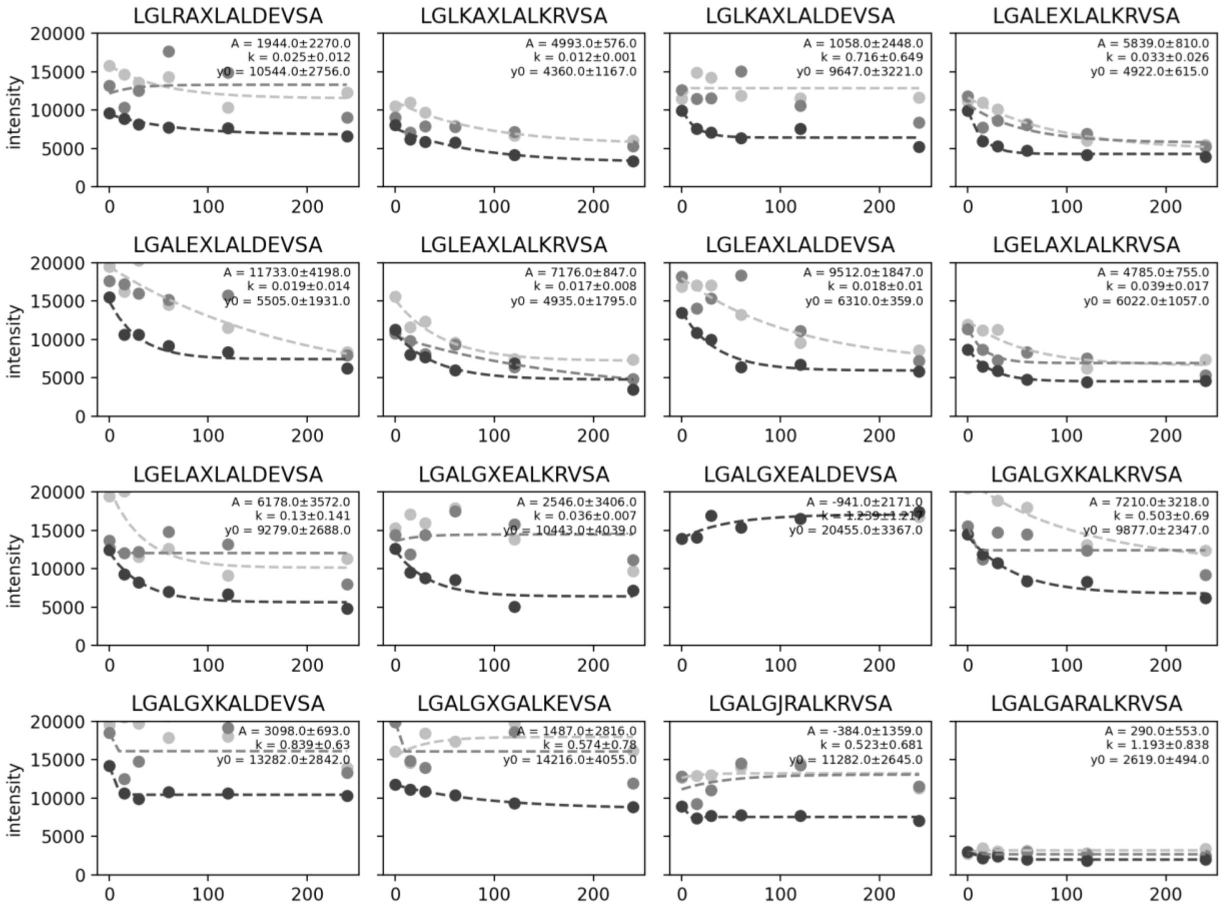


**Fig. S2 (continued).**

**A. (PP1)**

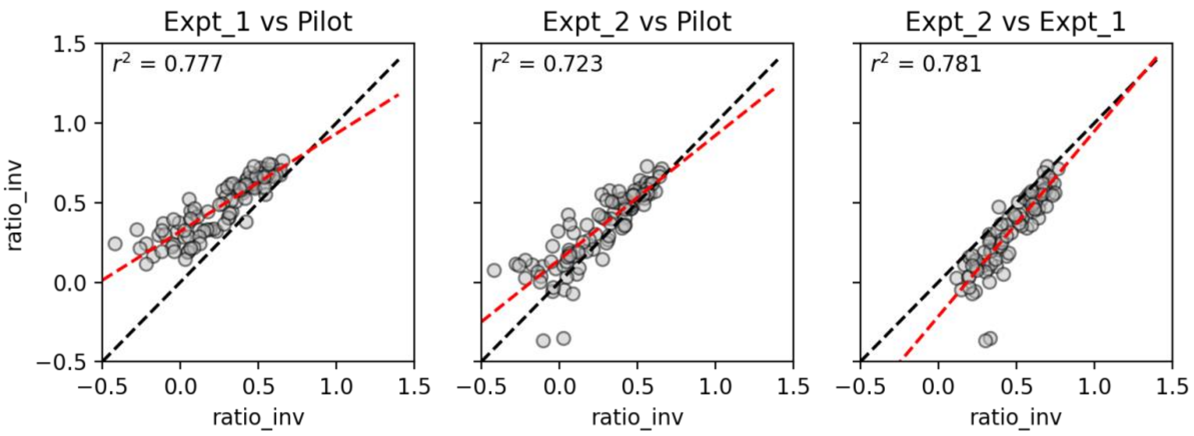


**B. (B55)**

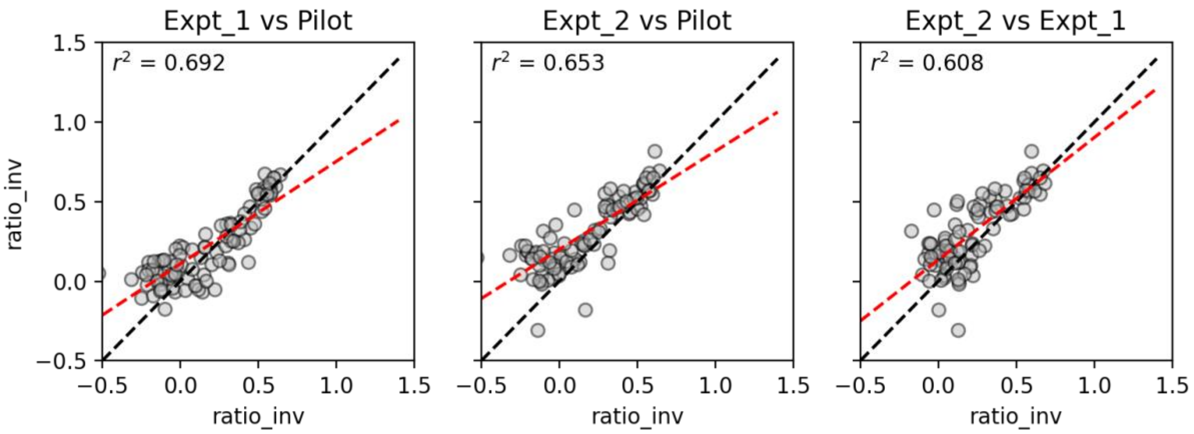


**Fig. S3.** Pairwise comparisons of dephosphorylation scores measured across 3 independent experimental replicates for PP1 **(A)** and PP2A-B55 (**B**). Each marker indicates the dephosphorylation score for a given peptide within a single experiment. Dashed black line signifies the identity line; red dashed line shows a linear regression; annotation specifies the Pearson correlation coefficient.

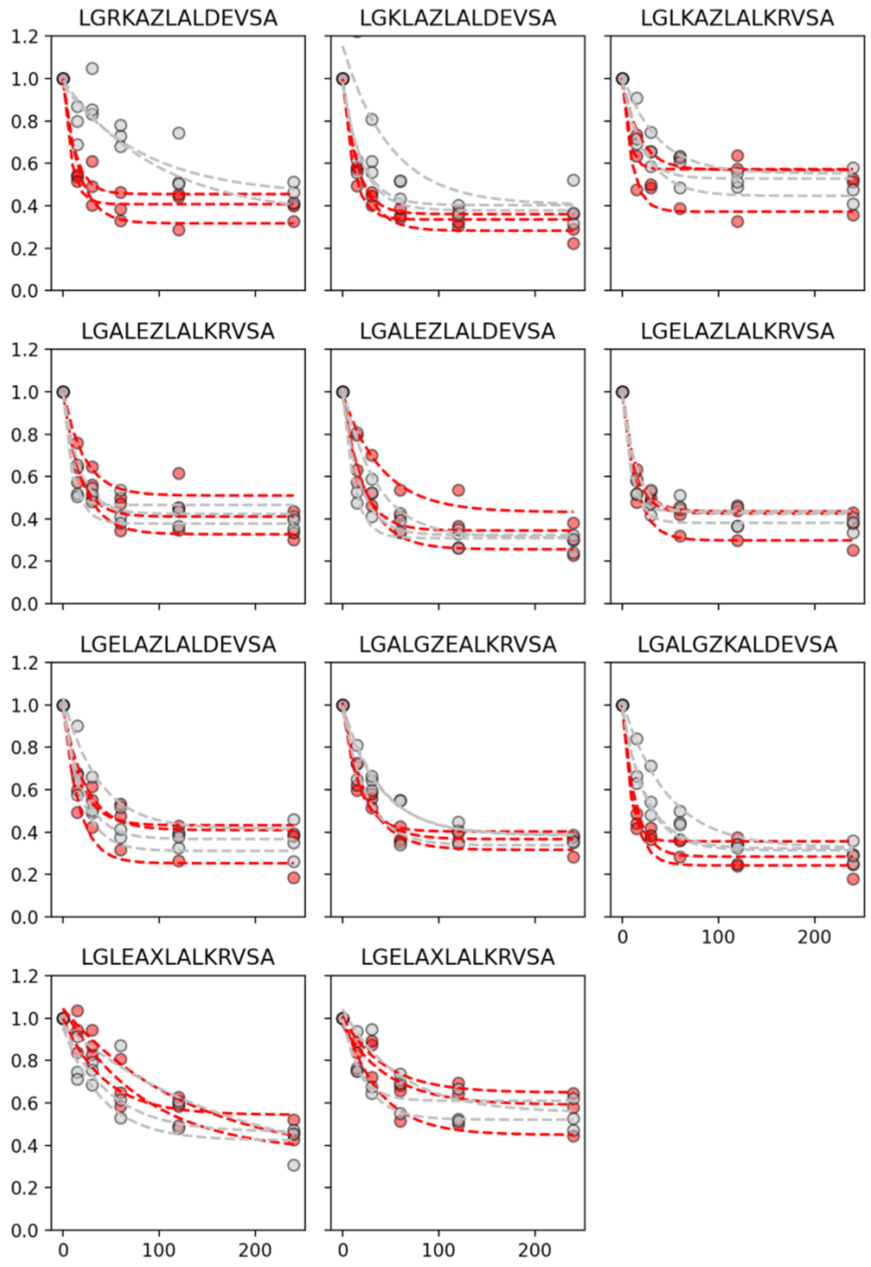


**Fig. S4.** Example progress curves showing measured DyLight 650 fluorescence (normalized to starting intensity) over time for PP1 (red) and B55 (grey) proteins for 11 peptides. Dashed lines indicate single exponential fit and highlight differences in relative kinetics of dephosphorylation across the 2 proteins.

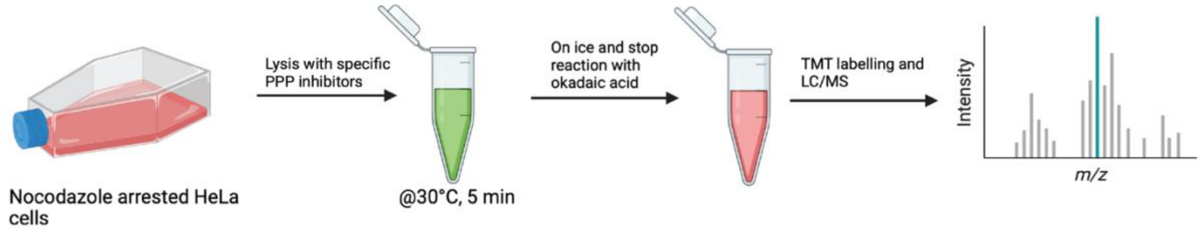


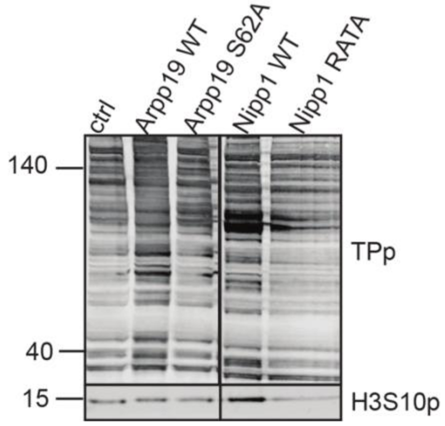


**Fig. S5.** Experimental setup of PPP inhibition in mitotic lysate and westernblot of total cell extract treated as indicated and probed for total phosphor TP and H3S10p.

A

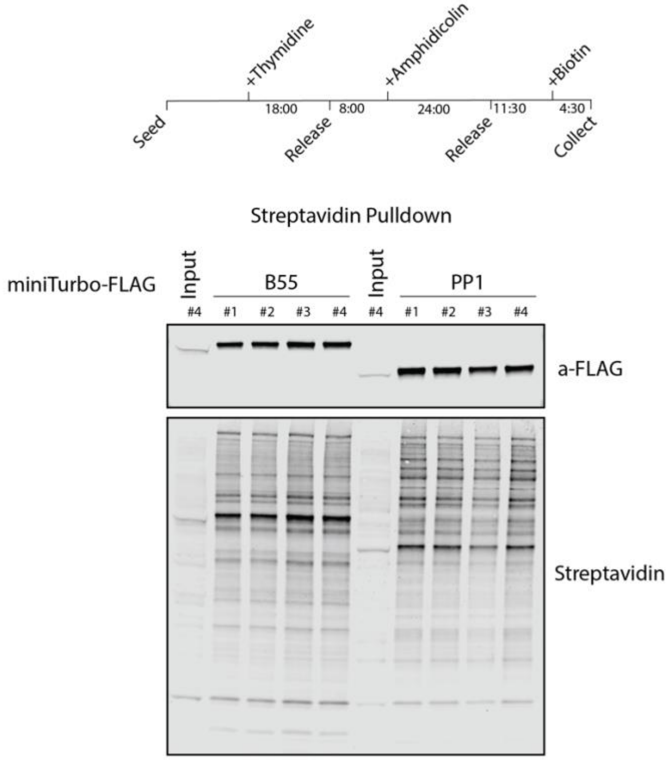


B

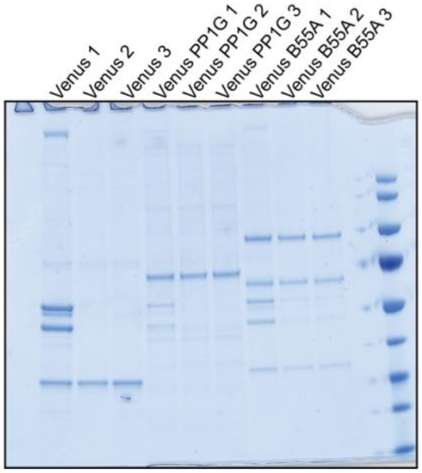


**Fig. S6.** Interactomes for PP1 and PP2A-B55. A) Synchronisation protocol used for miniTurboID experiments and analysis of samples by westernblot. B) Affinity purified YFP-tagged samples analysed by coomassie gel.

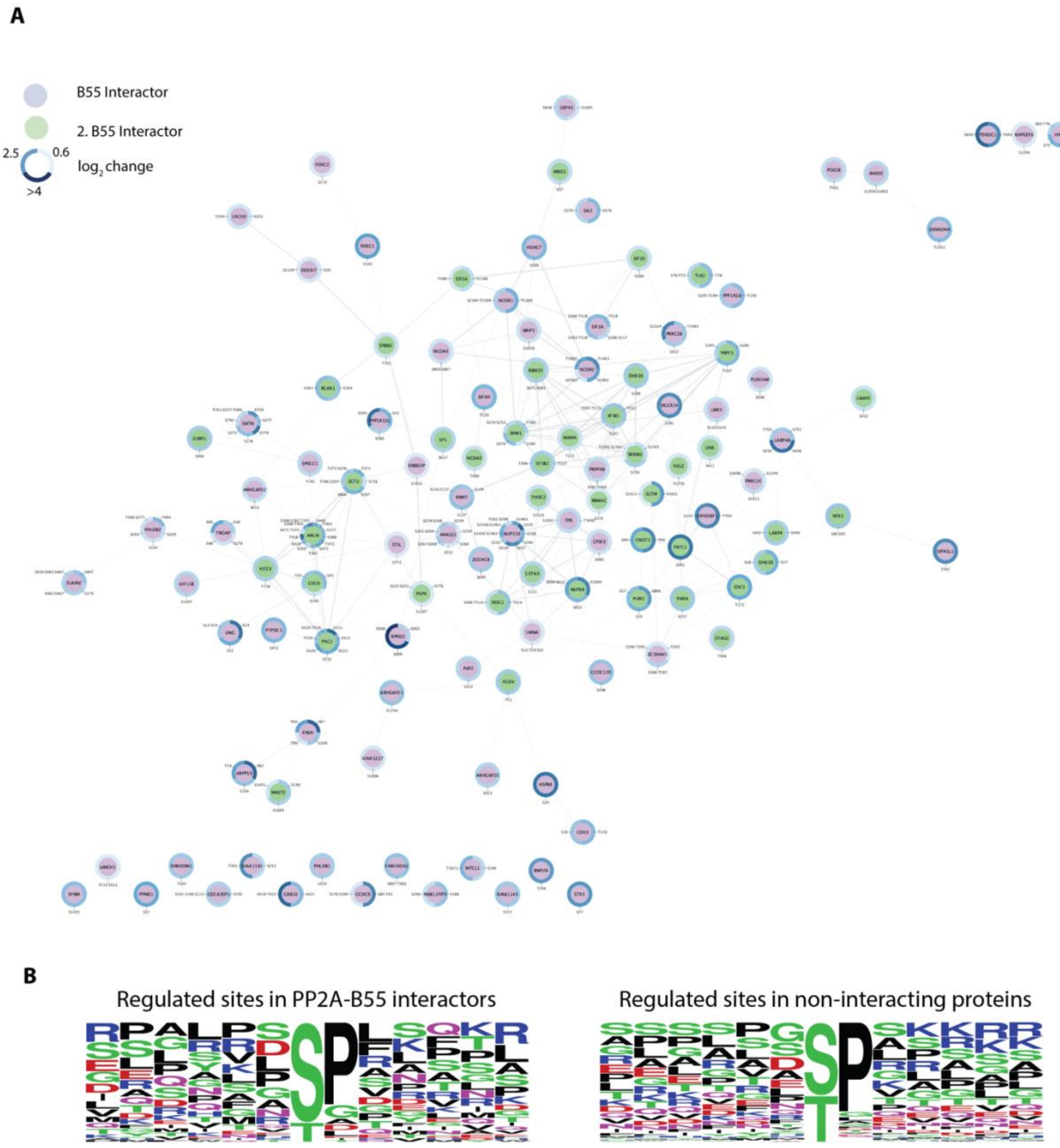


**Fig. S7.** A) B55 dephosphorylation network obtained by integrating interactors and mapped regulated sites. B) Analysis of regulated sites in PP2A-B55 interactors and in non-interacting proteins.

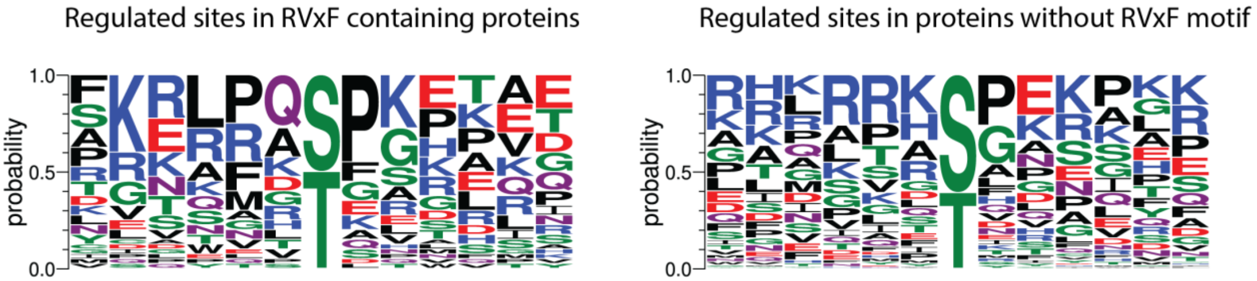


**Fig. S8.** Analysis of phosphorylation sites regulated by PP1 and whether the sites where present in proteins with or without an RVxF motif.

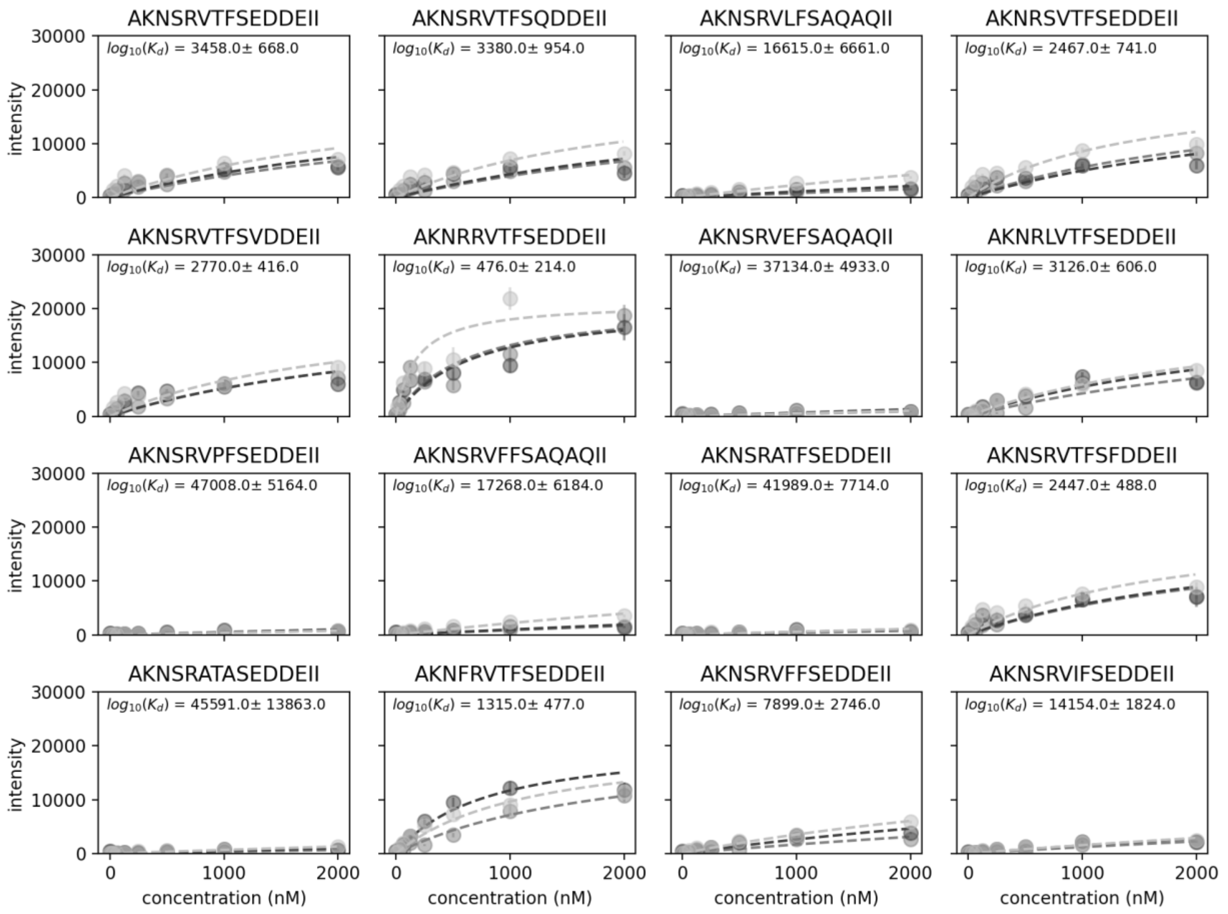


**Fig. S9.** MRBLE-pep concentration-dependent peptide binding curves for PP1. Each shade (markers and fits) indicates a different experimental replicate; dashed lines indicate a Langmuir isotherm fit ($y = y_{max}*[PP1]/([PP1] + K_d)$); annotated values are the mean ± standard deviation across 3 independent experimental replicates.

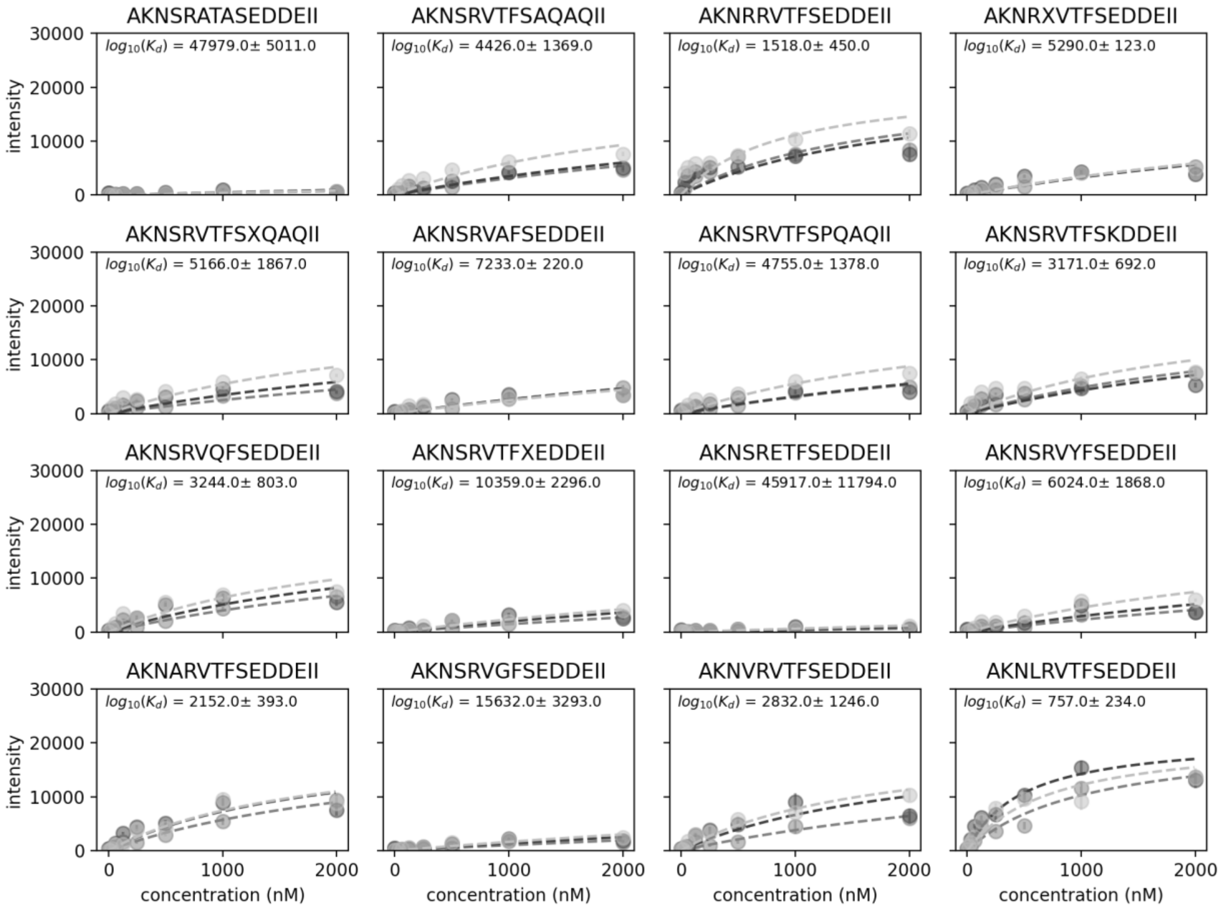


**Fig. S9 (continued).**

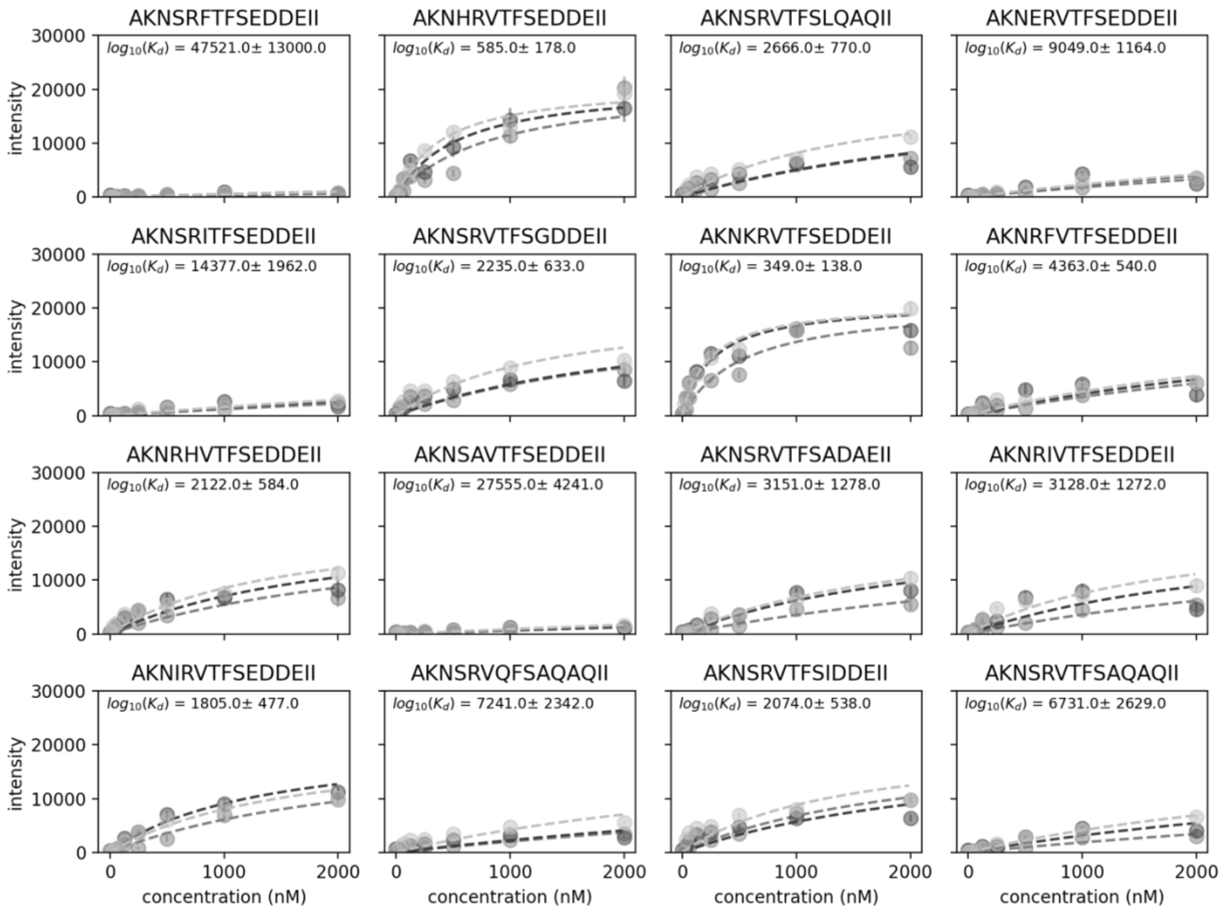


**Fig. S9 (continued).**

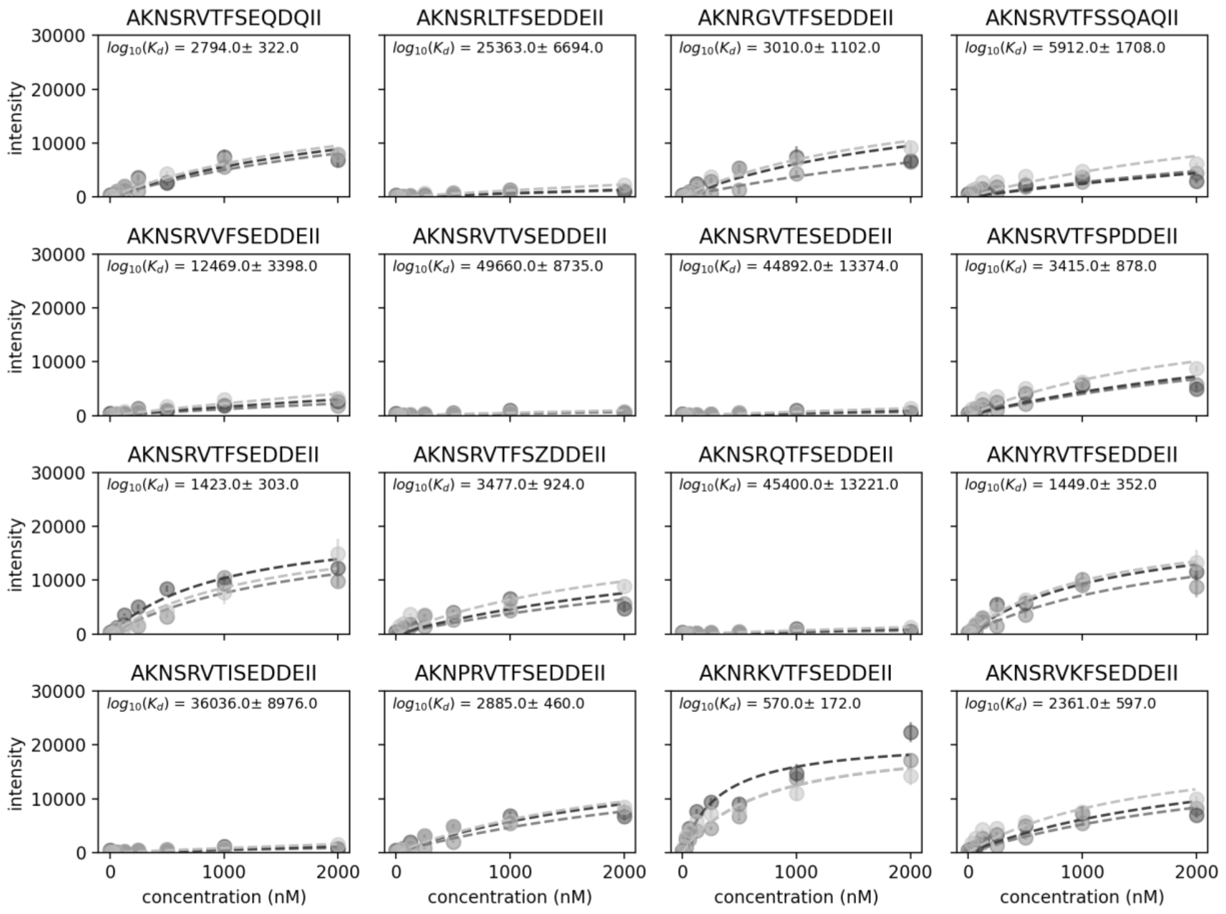


**Fig. S9 (continued).**

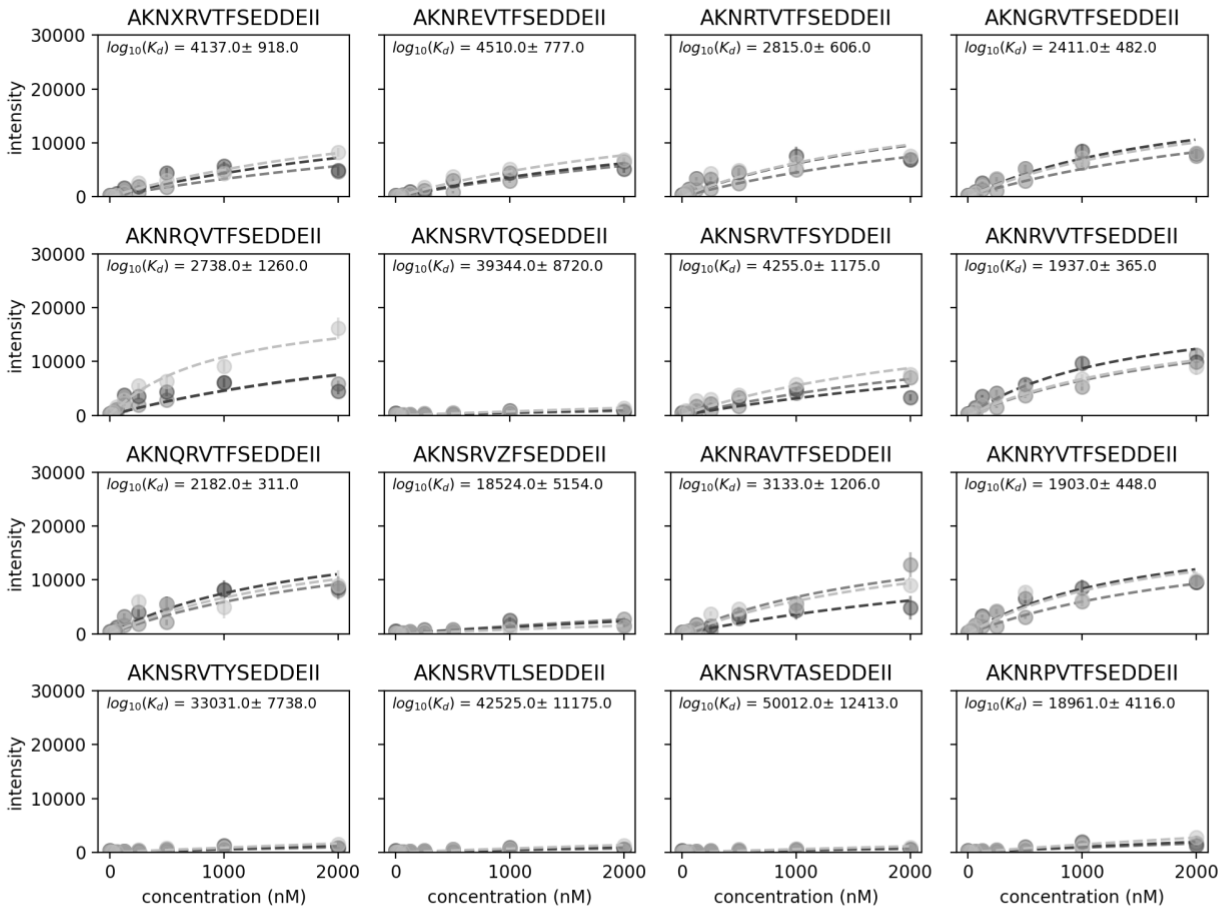


**Fig. S9 (continued).**

A

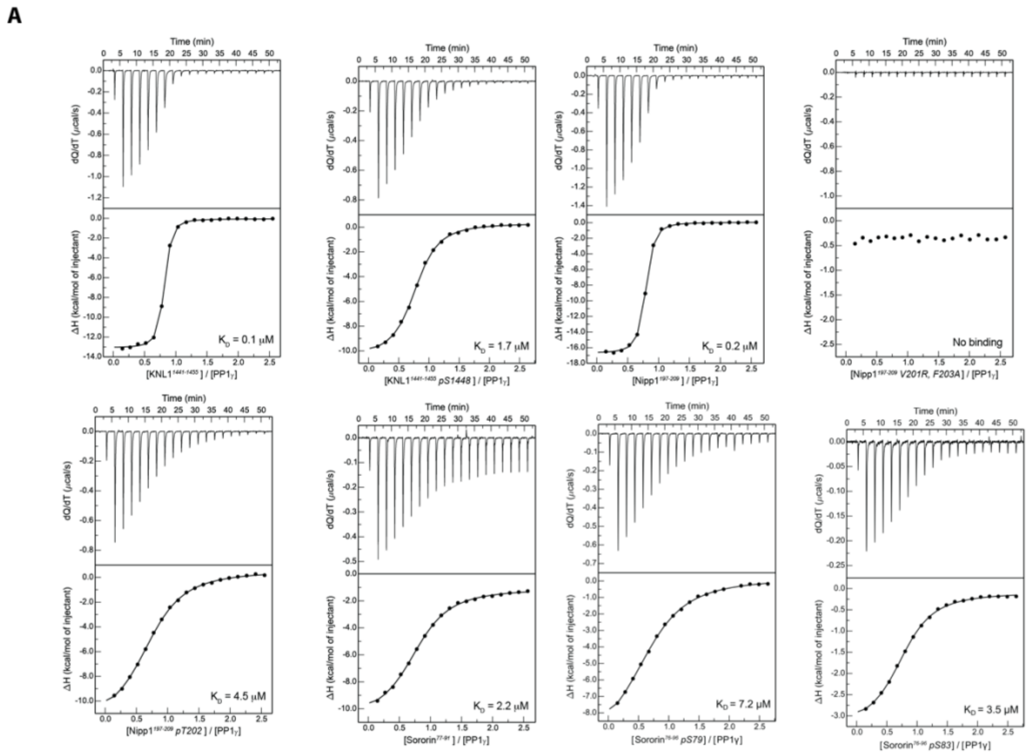


B

*Impact of RVxF motif phosphorylation on PP1 binding*

| PP1γ complex | Peptide sequence | $K_D$ (μM) |
|---|---|---|
| PP1γ/Nipp1$^{197-209}$ | AKNSRVTFSEDDEIW | 0.19 ± 0.01 |
| PP1γ/Nipp1$^{197-209}$ pT202 | AKNSRV(pT)FSEDDEIW | 4.51 ± 0.20 |
| PP1γ/Nipp1$^{197-209}$ V201R, F203A | AKNSRATASEDDEIW | No binding |
| PP1γ/Nipp1$^{197-209}$ V201R, pT202, F203A | AKNSRA(pT)ASEDDEIW | No binding |
| PP1γ/KNL1$^{1441-1455}$ | KLNSKRVSFKLPKDQW | 0.12 ± 0.01 |
| PP1γ/KNL1$^{1441-1455}$ pS1448 | KLNSKRV(pS)FKLPKDQW | 1.73 ± 0.01 |
| PP1γ/Sororin$^{77-91}$ pS83 | RRSPRI(pS)FFLEKENEW | 1.36 ± 0.01 |
| PP1γ/Sororin$^{76-96}$ | PRRSPRISFFLEKENEPPGREW | 2.53 ± 0.11 |
| PP1γ/Sororin$^{76-96}$ pS79 | PRR(pS)PRISFFLEKENEPPGREW | 7.24 ± 0.40 |
| PP1γ/Sororin$^{76-96}$ pS83 | PRRSPRI(pS)FFLEKENEPPGREW | 3.46 ± 0.18 |

C

*PP1 dependent phosphosites in RVxF motif*

| Protein | Modsite | RVXF | Position |
|---|---|---|---|
| NOC3L | S116 | GQRVSFL | 103, 110 |
| CDCA5 | S75, S83 | SPRISFF | 78, 85 |
| PGM2 | S165 | GYKVYWD | 172, 179 |
| ARHGH | S383 | VAKVSFP | 371, 378 |
| NOL10 | S475 | RFKVMFE | 483, 490 |
| SCAPE | S199 | ARRSLNFG | 196, 204 |
| FRM4B | S940 | SQRCLGFA | 925, 933 |
| TACC3 | T59 | AMKVTFQ | 51, 58 |
| KI67 | S507 | KRRRVSFG | 501, 509 |
| RBP2 | T1944 | NGRGVIFG | 1931, 1939 |
| RL24 | T83 | NPRQINWT | 71, 79 |
| PP1R8 | S199, T202, S204 | NSRVTFS | 197, 204 |
| PGM2L | S175 | GYKVYWE | 182, 189 |
| RIF1 | S2205 | KVRRVSFA | 2199, 2207 |
| CASC5 | S60, S1448 | NSRRVSFA - NSKRVSFK | 54,62 - 1442, 1450 |

D

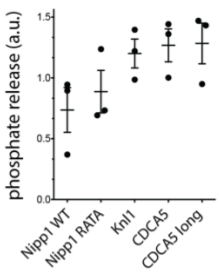


**Fig. S10.** A) ITC curves for the indicated peptides using PP1γ. B) Table of Kd values for the indicated peptides. C) PP1 regulated sites identified in our mitotic exit screen that resides in RVxF sequences. D) Dephosphorylation of RVxF model peptides by PP1.
